# FateDriver: Weakly supervised dual-view graph learning for transition-specific prioritization of candidate regulators in cellular reprogramming

**DOI:** 10.64898/2026.10.27.762150

**Authors:** Jieun Sung, Kyuwon Son, Hyuk-Jin Cha, Wankyu Kim, Haeseung Lee

**Affiliations:** Department of Life Science, Ewha Womans University, Seoul, Republic of Korea; Department of Pharmacy, College of Pharmacy, Pusan National University, Busan, Republic of Korea; College of Pharmacy and Research Institute of Pharmaceutical Sciences, Seoul National University, Seoul, Republic of Korea

**Author notes:** These authors contributed equally (Jieun Sung, Kyuwon Son). Correspondence (W.K.) and (H.L.).

**Keywords:** cellular reprogramming, single-cell RNA sequencing, graph neural network, weakly supervised learning, positive-unlabeled learning, cross-attention, induced pluripotency, cell-fate transition

## Abstract

Cellular reprogramming supports regenerative medicine, yet the endogenous regulators of a specified source-to-target transition remain incompletely defined. Existing computational methods rank candidate regulators from network topology, trajectory dynamics, or cell-state labels, leaving unused the factors experimentally delivered to induce the transition. We present Fat-eDriver, a weakly supervised dual-view graph neural network that jointly represents cells and genes from single-cell transcriptomes, treats the delivered factors as sparse gene-level labels, and ranks the remaining genes for a specified transition under a positive–unlabeled formulation. Across two OSKM datasets and one GMT dataset, FateDriver ranked first or second among six methods in recovering externally curated reference genes, exceeded a supervision-matched non-graph model despite weaker co-expression with the delivered factors, and achieved the highest recovery of protein targets of pluripotency-inducing chemical cocktails in both OSKM datasets. Holding either the source or the endpoint fixed yielded distinct candidate programs, including a *DDX58*–interferon axis associated with cells lacking endogenous pluripotency markers. Applying the same framework to candidate source–target pairs prioritized distinct candidate origins for two *in vivo* partially reprogrammed intestinal populations. FateDriver converts sparse experimental knowledge into transition-specific hypotheses for regulator prioritization from single-cell trajectories.

## INTRODUCTION

Cellular reprogramming redirects cell identity through defined transcription factors^1^ and provides a basis for regenerative medicine and disease modeling^2^. Related alterations in cell identity also contribute to disease progression in diverse tissues, including cancer^3–6^. The OSKM factors — *OCT4* (*POU5F1*), *SOX2, KLF4* and *MYC* — induce pluripotency and cellular plasticity^7,8^, whereas GMT factors — *Gata4, Mef2c*, and *Tbx5* — directly convert fibroblasts toward cardiomyocyte-like states^9^. These delivered factors initiate reprogramming but do not fully determine its progression or outcome. Cell-intrinsic regulatory systems shape reprogramming efficiency, trajectory, and cell-fate outcome^10–12^. At single-cell resolution, cells exposed to the same factors follow heterogeneous trajectories and reach distinct endpoints^13–15^. Single-cell RNA sequencing (scRNA-seq) resolves these trajectories and identifies genes whose expression changes during reprogramming. However, trajectory-associated expression alone does not distinguish genes that regulate the transition from downstream responses or markers of the resulting state. Identifying candidate regulators therefore remains a central challenge, particularly because the relevant genes may depend on both the source and target populations.

Existing computational methods infer candidate regulators from regulatory network topology, trajectory dynamics, or perturbations of cell-state representations^16,17^. CEFCON and SwitchTFI use prior gene regulatory networks and rank regulators through network control or centrality^18,19^; TENET reconstructs regulatory interactions along pseudotime and ranks genes by their down-stream connectivity^20^; and Fatecode ranks genes by perturbing a learned representation of cell state^21^. These approaches use regulatory structure, expression dynamics, or cell-state labels, but do not use the factors experimentally delivered to induce the transition as gene-level supervision. Network-based methods may additionally restrict the set of rankable genes to those represented in the regulatory prior or annotated as transcription factors, and topology-based scores may favor highly connected genes. Thus, existing approaches leave unused a small but experimentally grounded source of gene-level information already available in reprogramming experiments.

Using delivered factors as gene-level supervision presents two specific challenges. First, supervision is extremely sparse: only three or four delivered factors are known positives in each reprogramming system, whereas the remaining genes are untested rather than confirmed negatives. Treating all other genes as negatives could therefore penalize uncharacterized regulators. The task is more naturally formulated as positive–unlabeled-aware ranking than as binary classification^22,23^. Second, regulator relevance is expected to depend on both the starting and target states. Different source states may use distinct regulatory programs to reach the same target. Conversely, transitions from the same source toward different target states may involve distinct regulatory programs. Candidate regulators should therefore be prioritized for a specified source-to-target transition rather than inferred solely from their association with the target state.

Here, we present FateDriver, a weakly supervised dual-view graph neural network for transition-specific regulator prioritization from single-cell transcriptomes. FateDriver uses the transcription factors delivered in each experiment as sparse gene-level positive labels and ranks the remaining genes within a positive–unlabeled-aware framework conditioned on a specified source-to-target transition. Delivered factors form the highest-ranking tier, their annotated targets form an intermediate tier, and the remaining genes are treated as unlabeled. FateDriver jointly represents cells and genes while using prior regulatory information to inform gene representations without restricting scoring to genes represented in the prior. We benchmarked FateDriver against four published regulator-inference methods and a supervision-matched non-graph multi-layer perceptron (MLP) trained on the same single-cell data, pseudotime, and gene-level labels across OSKM- and GMT-induced reprogramming datasets. We then tested whether the rankings changed when either the source or endpoint was varied and extended the same framework to compare alternative candidate sources for two intestinal populations partially reprogrammed *in vivo*.

## RESULTS

### Overview of the FateDriver framework

FateDriver prioritizes candidate regulators for a specified cellular reprogramming transition (Figure 1A). The model takes three main inputs: single-cell gene-expression profiles of source and target cell populations together with an inferred ordering of cells along the reprogramming trajectory, or pseudotime, to represent the source-to-target transition. Its two graph encoders place cells and genes in a shared latent space, allowing each gene to be evaluated relative to the cellular transition. The identities of the transcription factors delivered in the experiment are also provided as input and serve as sparse positive labels. FateDriver then assigns a transition-specific score to each retained gene according to its learned relationship with the source, target, and transition. A higher score indicates greater predicted relevance to the specified source-to-target transition.

**Figure 1.**
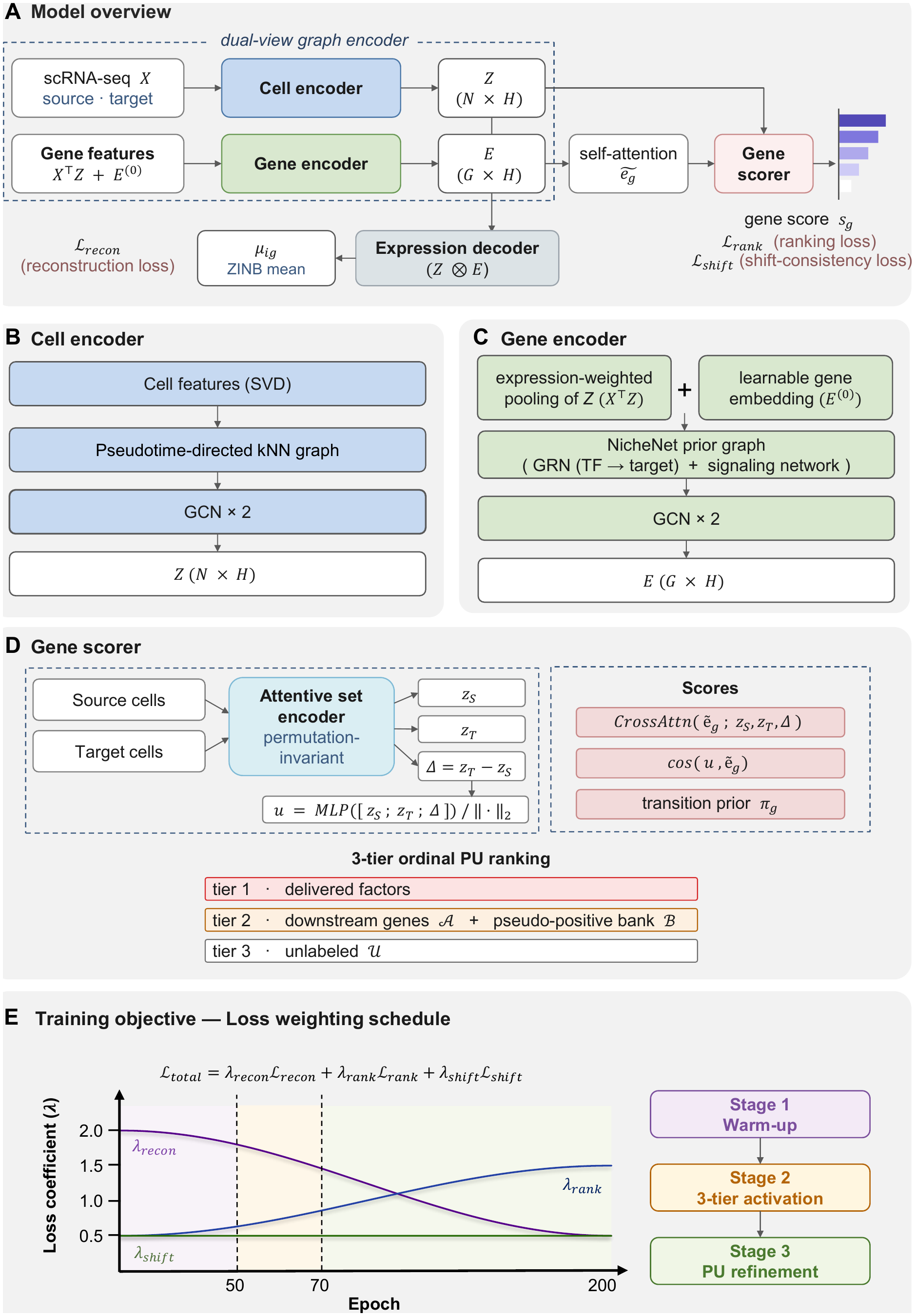
Overview of the FateDriver framework. (A) FateDriver jointly learns cell and gene embeddings to predict candidate regulators of a specified source-to-target transition. The expression decoder reconstructs observed counts from these embeddings, while the gene scorer assigns transition-specific gene scores. (B) The cell encoder applies two graph convolutional layers to expression features on a nearest-neighbor graph directed forward in pseudotime. (C) The gene encoder combines expression-weighted cell embeddings with learnable gene embeddings, then refines them over the NicheNet regulatory and signaling graph. (D) Source and target cell embeddings are pooled separately, and their difference defines the transition. Gene scores combine cross-attention, similarity, and transitionprior terms. Ranking supervision prioritizes delivered factors above an intermediate tier of annotated downstream targets and the pseudo-positive bank, followed by unlabeled genes. (E) Training combines reconstruction (ℒ_recon_), ranking (ℒ_rank_), and shift-consistency (ℒ_shift_) losses. Cosine interpolation decreases the reconstruction weight and increases the ranking weight, while the shift-consistency weight remains constant. The ranking curriculum introduces the annotated-target tier in stage 2 and progressively relaxes penalties involving unlabeled genes during stages 2 and 3. Parameter values are provided in STAR Methods, *Total loss and training schedule*.

FateDriver comprises four jointly trained modules: a cell encoder, a gene encoder, an expression decoder, and a gene-scoring head. The cell encoder maps cells into the shared space using their expression profiles and pseudotime ordering (Figure 1B). Cells with similar expression profiles are first connected in a nearest-neighbor graph. Only edges directed from earlier to later pseudotime are retained. A graph convolutional network then aggregates information from connected cells to generate the cell embeddings. The gene encoder maps genes into the same latent space as the cells (Figure 1C). For each gene, the encoder aggregates the embeddings of cells expressing that gene, weighted by its expression level, and combines the aggregated cell information with a gene-specific embedding learned during model training. A second graph convolutional network refines these representations using regulatory and signaling interactions from NicheNet^24^. Because the initial gene representations are constructed independently of network membership, genes absent from the prior graph can still be represented and scored. The expression decoder reconstructs the observed gene counts from the cell and gene embeddings, thereby anchoring the learned representations to the expression data; its output does not directly enter the gene score. The gene-scoring head evaluates each gene relative to the source population, target population, and source-to-target transition (Figure 1D). Self-attention first updates each gene representation using information from the other genes. The gene-scoring head then summarizes the source and target cell populations separately, and the difference between these summaries defines the transition vector from the source state to the target state. For each gene, the final score combines three complementary components: a cross-attention term that weights the source, target, and transition representations for that gene; a similarity term that measures cosine similarity between the gene representation and a learned scoring direction derived from the three cellular representations; and a transition-informed feature term based on gene-level expression change, pseudotime association, and regulatory-network connectivity.

Only three or four delivered factors are available as positive labels in each reprogramming system, whereas the remaining genes are unlabeled rather than confirmed negatives. Fat-eDriver therefore formulates the task as positive–unlabeled–aware ranking rather than binary classification^22^. Delivered factors form the highest tier, their annotated downstream targets form an intermediate tier, and all remaining genes are treated as unlabeled. The unlabeled group may contain unknown regulators and is therefore not treated as a set of confirmed negative genes. Genes that repeatedly receive high scores and remain close to labeled genes in the shared space can be moved into a pseudo-positive bank within the intermediate tier (STAR Methods, *Three-tier ranking objective*).

The four modules are trained jointly using three objectives (Figure 1E). First, the reconstruction loss (ℒ_recon_) trains the cell and gene embeddings to reproduce observed expression. Second, the ranking loss (ℒ_rank_) imposes the three-tier ordering. Third, the shift-consistency loss (ℒ _shift_) encourages highly ranked genes to align with the source-to-target transition. Training initially emphasizes reconstruction and gradually places more weight on gene ranking. Together, these components allow FateDriver to use sparse factor labels while ranking genes across the retained transcriptome (STAR Methods, *Total loss and training schedule*).

### FateDriver recovers external reprogramming benchmarks beyond factor co-expression

We evaluated FateDriver using three scRNA-seq datasets: two human OSKM-induced pluripotency datasets, GSE297234^25^ and GSE242423, and one mouse GMT-induced cardiac reprogramming dataset, GSE98567^26^ (Figure S1). We compared FateDriver with four published methods: CEFCON^18^, TENET^20^, Fatecode^21^, and SwitchTFI^19^. These methods use network structure, pseudotime dynamics, or cell-state labels, but they do not use the experimentally delivered factors as gene-level training labels. We therefore also included a non-graph MLP trained using the same single-cell data, pseudotime, delivered-factor labels, and annotated-target labels as FateDriver. The MLP represents each gene directly from its expression profile and pseudo-time values but does not construct graph-based cell and gene representations. This comparison was designed to assess whether FateDriver provided information beyond a simpler model given the same underlying data and supervision. All methods were evaluated against a common 3,000-gene universe (highly variable genes for each dataset), although CEFCON, TENET, and SwitchTFI returned partial rankings of 185–2,453 genes, reflecting the restricted gene coverage of their regulatory-network-based scoring (Figure 2).

**Figure 2.**
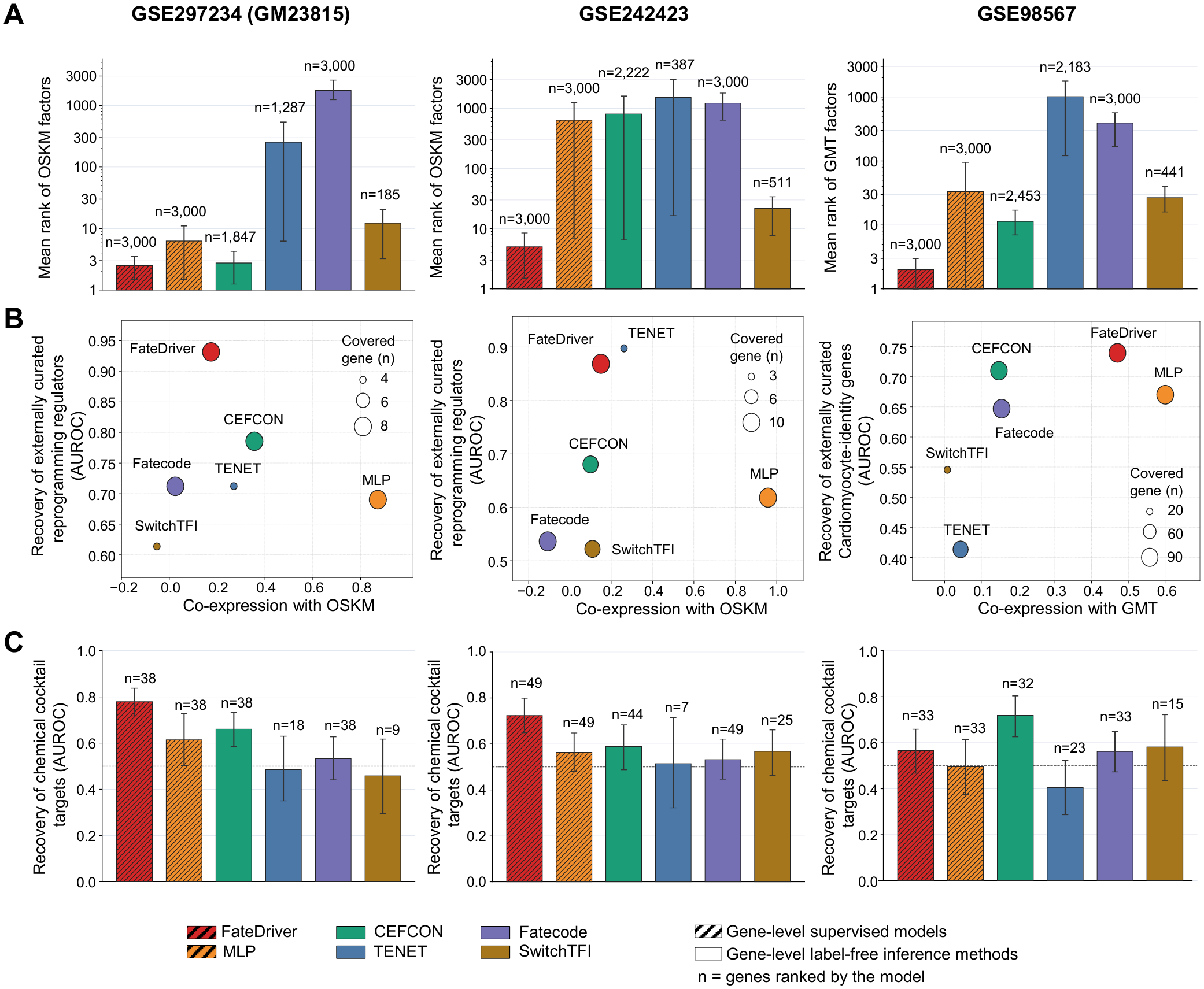
Benchmark of regulator prioritization across three reprogramming datasets. Columns show, from left to right, GSE297234 (donor GM23815; OSKM), GSE242423 (OSKM), and GSE98567 (GMT). (A) Mean rank of the delivered factors on a logarithmic axis; lower values indicate better ranks. Error bars show 95% gene-level bootstrap confidence intervals. A factor absent from a method’s ranking was assigned rank 3,000. Labels *n* indicate the total number of genes ranked by each method. (B) Co-expression with the delivered-factor module (*x* axis) versus recovery of a reference gene set (*y* axis). The reference set was externally curated reprogramming regulators for GSE297234 and GSE242423, and cardiomyocyte-identity genes for GSE98567. Co-expression is the mean Pearson correlation of the top-100 genes with the module along pseudotime. Recovery is the area under the receiver operating characteristic curve (AUROC) calculated from each method’s returned ranking. Delivered factors were excluded before both analyses. Marker size indicates the number of reference genes covered by each ranking. Marker sizes are scaled separately within each panel, with counts shown in the size key. (C) Recovery AUROC for the protein targets of reprogramming compounds: 262 targets of pluripotency-inducing compounds for GSE297234 and GSE242423, and 130 targets of cardiac-reprogramming compounds for GSE98567. Error bars show 95% bootstrap confidence intervals. The dashed line marks chance-level recovery (AUROC = 0.5). Labels *n* indicate the number of target genes covered by each method’s evaluated ranking. In A and C, hatched bars denote FateDriver and the MLP, which used delivered-factor and annotated-target labels during training. Plain bars denote the four methods that did not use these regulator gene labels. Colors distinguish methods throughout.

We first examined the ranks assigned to the delivered factors. Because these genes were used as positive labels by FateDriver and MLP, this analysis assessed whether the models retained the positive labels rather than providing an independent measure of regulator recovery. FateDriver assigned mean ranks of 2.5, 5.0, and 2.0 in GSE297234, GSE242423, and GSE98567, respectively, with lower values indicating better ranks (Figure 2A; Table S1). The corresponding MLP ranks were 6.3, 632.8, and 33.3. We next evaluated recovery of external reference genes after excluding the delivered factors. For the two OSKM datasets, the reference set comprised experimentally supported enhancers and barriers of pluripotency reprogramming^27^. Only three experimentally supported cardiac regulator genes were present in the analyzed gene space of GSE98567^28^. We therefore used curated cardiomyocyte-identity gene sets for that dataset^29–35^. FateDriver achieved the highest recovery AUROC among the six methods in GSE297234 (0.93) and GSE98567 (0.74) and ranked second in GSE242423 (0.87) (Figure 2B). TENET achieved the highest AUROC in GSE242423 (0.90), but its ranking contained only three reference genes among 387 ranked genes, compared with ten reference genes in the complete FateDriver ranking. FateDriver also yielded higher recovery point estimates than the supervision-matched MLP in all three datasets. The corresponding MLP AUROCs were 0.69, 0.62, and 0.67. Removing reference genes that overlapped the annotated-target training tier did not alter FateDriver’s relative performance (Table S2). We then examined whether external reference-gene recovery could be explained by simple co-expression with the delivered factors. The top-100 genes selected by the MLP showed the strongest co-expression with the delivered factors in all three datasets, with correlations of 0.87, 0.96, and 0.60 in GSE297234, GSE242423, and GSE98567, respectively (Figure 2B). The corresponding correlations for FateDriver were lower, at 0.17, 0.15, and 0.47. Despite this weaker co-expression, FateDriver yielded higher external-set recovery than the MLP in each dataset. Thus, genes with expression patterns most similar to those of the delivered factors were not necessarily the genes that best recovered the external reference sets.

As a second external evaluation, we examined whether the rankings recovered protein targets of small-molecule cocktails reported to induce pluripotency or cardiac conversion^36–46^. These target sets represent alternative interventions toward related cell-state endpoints. FateDriver achieved the highest chemical-target recovery among the six methods in both OSKM datasets (Figure 2C). The AUROCs were 0.78 (95% CI, 0.72–0.84) in GSE297234 and 0.72 (95% CI, 0.65– 0.80) in GSE242423. CEFCON ranked second, with AUROCs of 0.66 and 0.59, respectively. Excluding the ten chemical-target genes that also occurred in the annotated-target training tier yielded similar FateDriver AUROCs of 0.78 and 0.71. In the cardiac dataset, FateDriver achieved an AUROC of 0.57 (95% CI, 0.47–0.66), compared with 0.72 (95% CI, 0.63–0.80) for CEFCON.

The methods also generated substantially different top-ranked gene sets. In GSE297234, 37– 67 genes in each top-100 set were absent from the top-100 sets of all other methods, with similar patterns in the other datasets (Figure S2A–C, left). Among top-ranked genes represented in the NicheNet graph, FateDriver selected genes with higher median out-degree than the MLP but lower median out-degree than CEFCON and SwitchTFI (Figure S2A–C, right; Table S3). These comparisons indicate that FateDriver did not simply reproduce either the genes most strongly correlated with the delivered factors or the most highly connected network genes.

### Source-state conditioning yields distinct gene rankings toward a common iPSC endpoint

FateDriver is designed to rank genes for a specified source-to-target transition. We tested whether the gene ranking depends on the source population when the target population is held constant. GSE181451 contains two transcriptionally distinct mouse embryonic fibroblast (MEF) states, MEF-1 and MEF-2, connected to a common iPSC endpoint^10^. MEF-1 occupied the earliest region of the inferred trajectory, whereas MEF-2 was positioned closer to iPSC (Figure 3A). Compared with MEF-1, MEF-2 showed lower expression of MEF-enriched^10^, epithelial-to-mesenchymal transition (EMT), and senescence gene sets, and higher expression of embryonic stem-cell (ESC)-core and E2F-target gene sets (Figure S3A).

**Figure 3.**
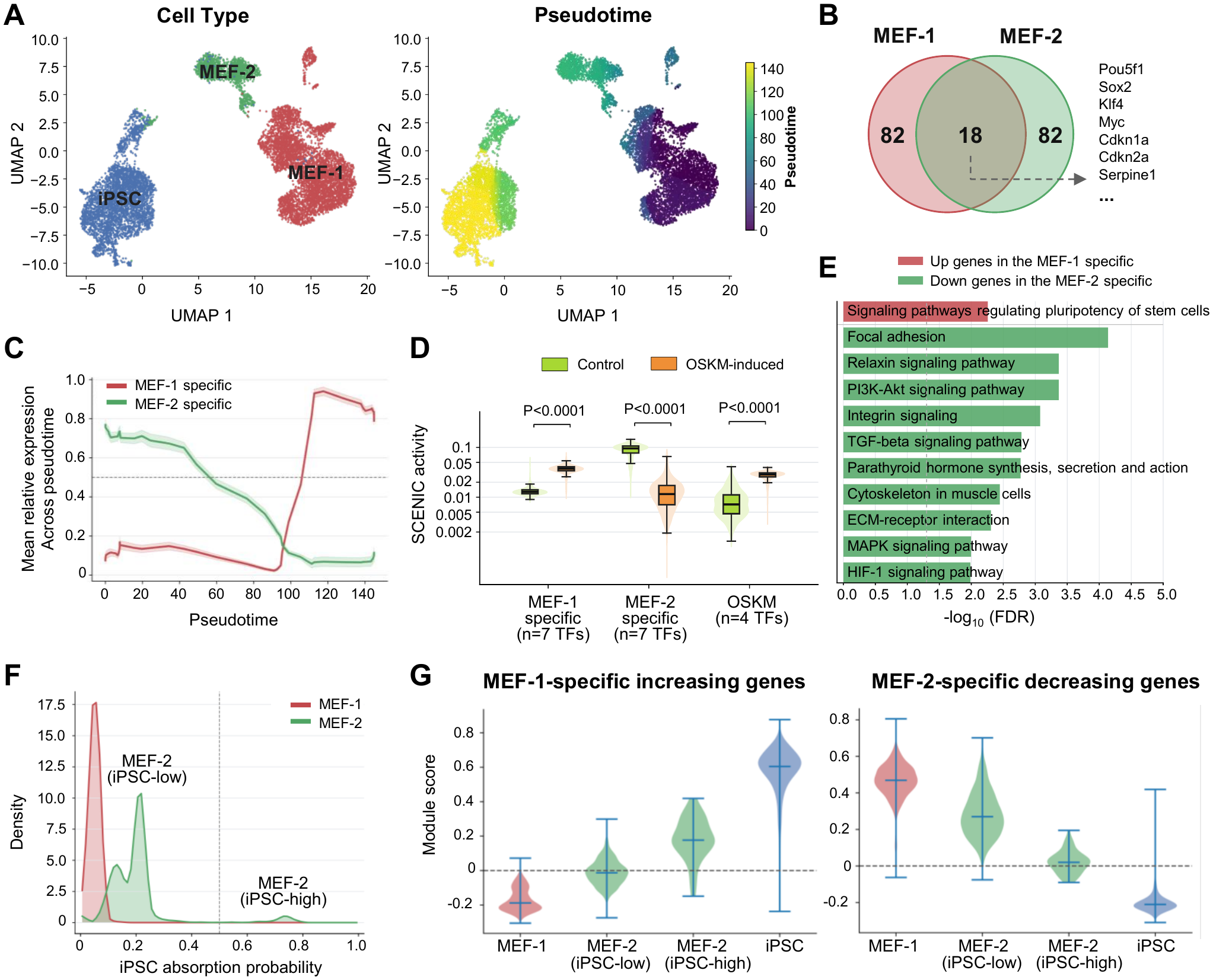
Source-state-dependent gene prioritization toward a common iPSC endpoint. (A) UMAP colored by cell state (left) and pseudotime (right). (B) Overlap of the top-100 genes prioritized for the MEF-1→ iPSC and MEF-2→ iPSC transitions. (C) Each gene’s smoothed mean expression was scaled to 0–1 across pseudotime bins. Curves show the average across genes in each source-specific set. Shading indicates the standard error across genes; the dashed line marks half-maximal expression. (D) Distributions of mean regulon activity for source-specific and shared transcription factors in control MEF and OSKM-induced iPSC cells, measured by pySCENIC AUCell. Boxes show medians and interquartile ranges; whiskers extend to the most extreme values within 1.5 interquartile ranges. *P* values are from two-sided Kolmogorov–Smirnov tests. (E) KEGG enrichment of MEF-1-specific genes and MEF-2-specific genes. Bar lengths indicate −log_10_(FDR). (F) Distributions of CellRank iPSC absorption probabilities in MEF-1 and MEF-2. The dashed line at 0.5 separates the MEF-2 iPSC-low and iPSC-high groups. (G) Per-cell module-score distributions for MEF-1-specific genes (left) and MEF-2-specific genes (right) across the four indicated cell groups. Horizontal marks show medians, capped lines span the minimum to maximum scores, and dashed lines mark zero.

FateDriver was applied separately to the MEF-1 →iPSC and MEF-2→ iPSC transitions using the same iPSC population as the target. The two top-100 rankings shared 18 genes, whereas 82 genes were specific to each source-state model (Figure 3B; Table S4). The shared set included all four delivered factors (*Pou5f1, Sox2, Klf4*, and *Myc*) and the senescence-associated reprogramming barriers *Cdkn1a, Cdkn2a*, and *Serpine1*^47,48^.

The two source-specific sets showed opposing patterns at both the gene-expression and inferred regulon-activity levels. Along pseudotime, 79 of the 82 MEF-1-specific genes increased toward iPSC, whereas 75 of the 82 MEF-2-specific genes decreased (Figure 3C). The transcription factors in each set had motif-supported regulons suitable for activity inference. Their median regulon activity increased from 0.013 to 0.037 for the MEF-1-specific factors (*Etv5, Klf2, Mybl2, Nanog, Neurod1, Zfp42*, and *Zic3*) and decreased from 0.095 to 0.012 for the MEF-2-specific factors (*Cebpb, Fhl2, Mxd4, Mxi1, Nfia, Nr2f2*, and *Prrx1*) (Figure 3D; STAR Methods, *Regulon activity* ). The shared OSKM factors also showed increased activity.

Pathway enrichment also differed between the two rankings. MEF-1-specific genes that increased along pseudotime were enriched for Signaling pathways regulating pluripotency of stem cells (FDR = 5.5× 10^*−*3^). MEF-2-specific genes that decreased along pseudotime were enriched for Focal adhesion (FDR = 7.3 ×10^*−*5^), TGF-*β* signaling (1.6 ×10^*−*3^), and ECM–receptor interaction (4.9× 10^*−*3^) (Figure 3E). EMT-signature expression in MEF-2 was lower than in MEF-1 but remained above iPSC levels (Figure S3A), consistent with progressive attenuation of mesenchymal programs during fibroblast reprogramming^49^. Differential-expression analyses of the two source-to-iPSC comparisons also showed corresponding pathway differences (Figure S3B–E).

We further examined whether expression of the source-specific gene sets tracked the inferred propensity of individual cells to reach the common iPSC endpoint. CellRank was used to estimate, for each cell, the probability of reaching the iPSC terminal state, referred to as the iPSC absorption probability^50^ (STAR Methods, *Fate-probability analysis using CellRank* ). The median probability was 0.050 in MEF-1 and 0.203 in MEF-2 (Figure 3F). MEF-2 cells were divided into low- and high-iPSC-probability groups using a CellRank probability threshold of 0.5 (*n* = 1,441 and *n* = 64, respectively). Within MEF-2, the high-iPSC-probability group showed higher expression of the MEF-1-specific increasing genes and lower expression of the MEF-2-specific decreasing genes than the low-iPSC-probability group (Figure 3G).

Thus, holding the iPSC target constant while changing the source state yielded largely distinct gene rankings that differed in expression direction, inferred regulon activity, pathway enrichment, and association with iPSC fate probability.

### Endpoint conditioning yields distinct gene rankings for partial and pluripote associated reprogramming states

We next tested whether FateDriver rankings depended on the target state when the source population was held constant. To address this question, we revisited GSE297234, which profiled OSKM-induced human fibroblasts at days 0, 3, 7, and 10 of reprogramming. In our initial benchmark (Figure 2), day-0 (D0) fibroblasts and the entire day-10 (D10) population were used as the source and operational iPSC-like target, respectively. As the original study identified a transcriptionally distinct partial-reprogramming state within this time course^25^, we integrated cells across the sampled time points and annotated clusters using published reprogramming signatures (Figure 4A,B; Figure S4A,B; Table S5). Cluster 4 showed a partial-reprogramming signature without a corresponding pluripotency signature and was therefore designated as the partial-reprogramming endpoint.

**Figure 4.**
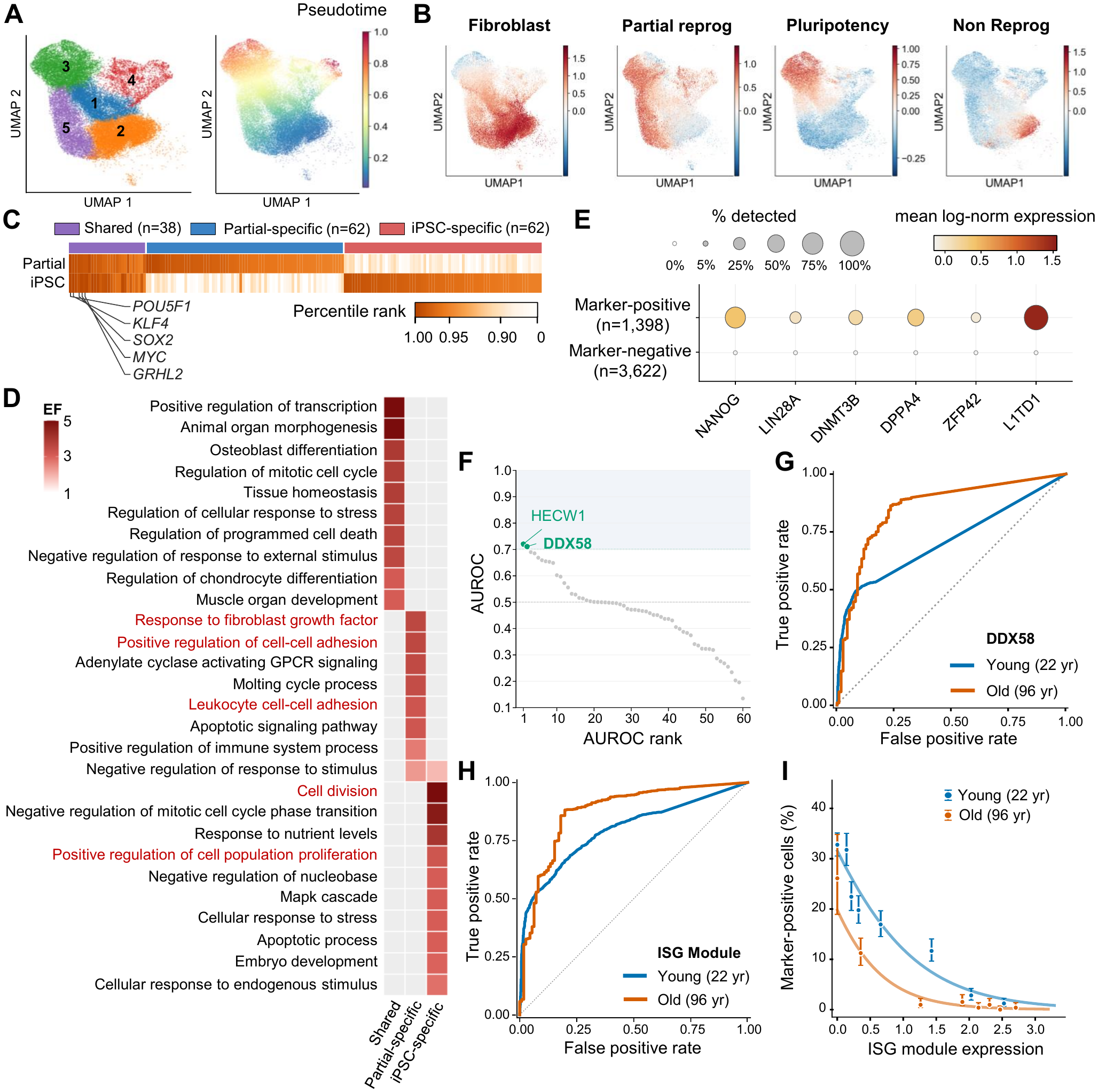
Endpoint-dependent gene prioritization and interferon-associated reprogramming outcomes. (A) UMAP of cells from the young donor (GM23815, 22 yr) from D0, D3, D7, and D10, colored by cell cluster (left) and pseudotime (right). (B) UMAPs showing fibroblast, partial-reprogramming, pluripotency, and non-reprogramming signature scores. (C) Heatmap of FateDriver rank percentiles for the fibroblast→ partial and fibroblast →iPSC-like transitions. The top-100 rankings contained 38 shared and 62 endpoint-specific genes each; higher percentiles indicate higher priority. (D) Gene Ontology biological-process enrichment of shared and endpoint-specific gene sets. Color indicates enrichment factor (EF). (E) Dot plot of six endogenous pluripotency markers in marker-positive and marker-negative D10 cells of the young donor. (F) AUROC values and corresponding ranks for 62 partial-specific genes in the young donor. Marker-negative cells were treated as the positive class; AUROC above 0.5 indicates higher gene expression in cells lacking detectable expression of the selected pluripotency markers. (G, H) ROC curves for *DDX58* expression (G) and interferon-stimulated gene (ISG) module score (H) distinguishing marker-negative from marker-positive D10 cells in young (GM23815, 22 yr) and old (GM00731, 96 yr) donors. (I) Percentage of marker-positive D10 cells across eight strata of an ISG module score. Points indicate observed percentages with Wilson 95% confidence intervals; curves represent sequencing-depth-adjusted logistic regression.

Using the same D0 fibroblast source, we generated separate FateDriver rankings for the fibroblast→ partial and fibroblast →iPSC-like transitions. The two top-100 rankings shared 38 genes, including all four delivered OSKM factors and the epithelial reprogramming regulator *GRHL2*, whereas 62 genes were specific to each endpoint (Figure 4C; Table S6). The endpoint-specific sets showed distinct functional enrichments (Figure 4D). Partial-specific genes were enriched for *response to fibroblast growth factor* (enrichment factor 3.6) and *positive regulation of cell–cell adhesion* (3.6), the latter consistent with mesenchymal-to-epithelial remodeling during fibroblast reprogramming^49^. In contrast, iPSC-like-specific genes were enriched for *cell division* (enrichment factor 5.3) and *positive regulation of cell population proliferation* (3.2).

A subset of D10 cells fell within the partial-reprogramming cluster (cluster 4; Figure S4A), indicating that the D10 population was heterogeneous in the reprogramming state. We therefore asked whether partial-specific genes marked D10 cells that had not activated endogenous pluripotency. D10 cells were classified by detection of six endogenous pluripotency markers (*NANOG, LIN28A, DNMT3B, DPPA4, ZFP42*, and *L1TD1*)^51,52^ (Figure 4E). Marker-positive cells comprised 19.1% of D10 cells in the young donor and 2.9% in the aged donor, consistent with the donor difference reported in the original study^25^. For each of the 62 partial-specific genes, we quantified by AUROC how well higher expression identified marker-negative cells (STAR Methods, *Day-10 reprogramming outcome and AUROC analysis*). Only *HECW1* and *DDX58* exceeded an AUROC of 0.70 (Figure 4F). Notably, only *DDX58* also exceeded 0.70 in the aged donor (GM00731, 96 years), where it discriminated marker-negative from marker-positive cells more strongly than in the young donor (AUROC, 0.851 versus 0.711) (Figure 4G). We therefore focused on *DDX58*, the only partial-specific candidate that preferentially marked D10 cells lacking endogenous pluripotency-marker expression in both donors.

*DDX58* encodes RIG-I, a cytosolic sensor of double-stranded RNA that activates type I interferon signaling^53^. We therefore tested whether downstream interferon-response genes captured this association more strongly than *DDX58* expression alone. A module score of four interferon-stimulated genes (ISGs; *ISG15, IFIT1, IFIT3*, and *IRF7* ) achieved higher AUROCs than *DDX58* alone in both the young (0.795) and the aged (0.873) donors (Figure 4H). Across eight ISG-score strata, the marker-positive fraction fell from 32.8% in the lowest stratum to 1.2% in the highest in the young donor, and from 26.1% to 0.4% in the aged donor (Figure 4I; Table S6). Because both marker detection and ISG scores can vary with per-cell sequencing depth, we adjusted for total transcript count. Higher ISG scores remained associated with lower odds of marker positivity in both donors (odds ratio, 0.35 in the young donor and 0.16 in the aged donor). ISG expression was therefore inversely associated with endogenous pluripotency-marker expression at day 10 in both donors, after accounting for sequencing depth.

Thus, conditioning on the partial endpoint prioritized *DDX58*, which was absent from the iPSC-like top-100 ranking and whose downstream interferon program was associated with the absence of endogenous pluripotency at day 10.

### FateDriver prioritizes candidate source populations for partially reprogramm intestinal cells

The preceding analyses ranked regulators for defined source–target pairs. We next examined whether FateDriver could prioritize candidate source populations when the target state was known but its cellular origin was uncertain. We reasoned that pairing a target with its true source should allow FateDriver to model the reprogramming transition more accurately, so that known reprogramming regulators would receive higher ranks than with an incorrect source.

To test this hypothesis, we applied FateDriver to GSE237173, a mouse intestinal dataset in which doxycycline (Dox) treatment induced OSKM expression *in vivo*, and annotated nine epithelial populations using markers from the original study^54^ (Figure S5A–C). Overall, Dox treatment increased partial-reprogramming scores^25^ and decreased pluripotency scores (Figure S5D). Among these populations, revival stem cell-like (revSC) and atrophy-induced villus epithelial cell-like (aVEC) cells expanded upon Dox treatment (Figure S5A,B), showed the largest Dox-induced increases in partial-reprogramming score (Figure 5A), and had the highest YAP and fetal signature scores (Figure S5E), consistent with the original study^54^. We therefore treated revSC and aVEC as two partially reprogrammed target populations. Trajectory analysis toward each of these two targets identified four candidate lineages, originating from crypt base columnar cells (CBC), transit-amplifying cells (TA), enterocytes (EC), and Paneth cells (PC) (Figure 5B).

**Figure 5.**
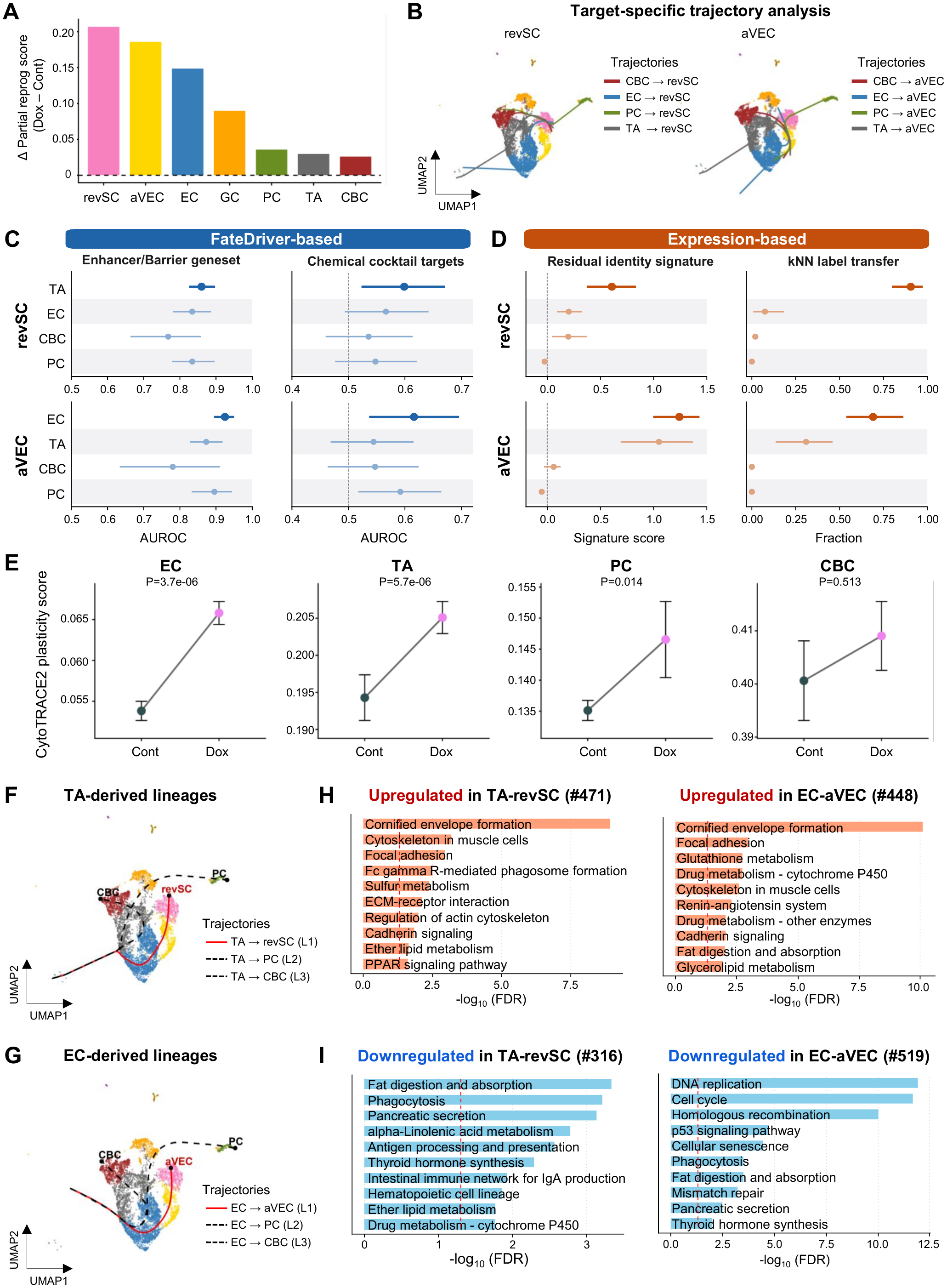
Prioritization of candidate source populations for partially reprogrammed intestinal cells. (A) Differences in partial reprogramming signature scores between doxycycline (Dox) and control (Cont) across intestinal cell types. Cell types with fewer than 50 cells in the Dox condition were excluded. (B) Candidate trajectories from four source populations (CBC, EC, PC, and TA) toward revSC (left) or aVEC (right), overlaid on UMAP embeddings. (C) Recovery AUROCs for two curated reference gene sets (enhancer/barrier reprogramming regulators and chemicalcocktail targets) using FateDriver rankings from each candidate–target pair. Error bars indicate 95% bootstrap confidence intervals from gene resampling; dashed lines mark AUROC = 0.5. (D) Expression-based assessment of candidate sources using residual source-identity signature scores (left) and k-nearest-neighbor (kNN) label transfer (right). Error bars indicate 95% intervals from cell resampling; the dashed line marks a signature score of zero. Top-ranked candidates are highlighted. (E) CytoTRACE2 plasticity scores for each candidate source population in Cont and Dox conditions. Data are mean± SEM; *P* values were calculated using one-sided Wilcoxon rank-sum tests. (F, G) Trajectories rooted at TA (F) or EC (G). Red lines indicate trajectories toward revSC or aVEC (L1), respectively; dashed black lines indicate alternative lineages (L2 and L3). (H, I) KEGG pathway enrichment of genes specifically upregulated (H) or downregulated (I) along the TA–revSC and EC–aVEC trajectories. Numbers in parentheses indicate the corresponding gene counts.

For each target, we trained FateDriver separately with each candidate source, keeping the target cells and the number of source cells fixed, and compared how well each model recovered the two external reference gene sets: curated reprogramming regulators and chemical-cocktail targets. We hypothesized that a more plausible source–target pairing would yield higher recovery of these reference genes. For revSC, the TA-based model achieved the highest AUROCs for both curated regulators (0.860) and chemical-cocktail targets (0.599). In contrast, the EC-based model ranked first for aVEC (0.925 and 0.616, respectively), followed by PC (0.895 and 0.592) (Figure 5C; Table S7). In paired bootstrap resampling of reference and non-reference genes, TA outperformed each alternative source for revSC in at least 84% of comparisons, and EC outperformed TA and CBC for aVEC in at least 97% of comparisons but exceeded PC in 80–90% (STAR Methods, *Candidate-source prioritization in GSE237173*). Gene-set recovery therefore favored TA as the source of revSC and EC as the source of aVEC, with EC less clearly separated from PC.

To test whether these predictions were supported independently of FateDriver, we examined candidate origins using two types of expression-based evidence: residual source identity in target cells and OSKM-induced plasticity in candidate sources. First, we assessed residual source identity with two separate measures: an identity-signature score for each candidate in target cells, and the fraction of target cells assigned to each candidate by nearest-neighbor label transfer from control cells (Figure 5D; STAR Methods, *Expression-based assessment of the candidate sources*). Both measures ranked TA first for revSC and EC first for aVEC. Second, we compared CytoTRACE2 developmental-potential scores^55^ between Dox-treated and control cells of each candidate. EC and TA showed the most significant Dox-associated increases, followed by PC, whereas CBC showed no significant change (Figure 5E). Residual source identity therefore supported TA as the source of revSC and EC, rather than PC, as the source of aVEC, and the plasticity readout ranked the same two cell types highest.

Finally, we characterized the two predicted transitions. Trajectory analysis rooted at each predicted source yielded three lineages: the first led to its partially reprogrammed target (TA→ revSC or EC →aVEC; L1), whereas the second and third led to PC (L2) and CBC (L3) from both sources (Figure 5F,G). We then identified genes differentially expressed in L1 relative to both L2 and L3 (Figure S5F,G). Genes upregulated along TA →revSC (n = 471) and EC→ aVEC (n = 448) were both enriched for focal adhesion, cytoskeletal, and cadherin signaling pathways (Figure 5H), consistent with the adhesion remodeling that accompanies early reprogramming^49,56^. In contrast, the downregulated programs diverged; genes downregulated along TA→ revSC (n = 316) were enriched for mature intestinal epithelial functions, including fat digestion and absorption, pancreatic secretion, and antigen processing and presentation (Figure 5I). Genes downregulated along EC →aVEC (n = 519) were enriched for DNA replication, cell cycle, homologous recombination, cellular senescence, and p53 signaling; the reduction in p53 and senescence signaling agrees with a previous report that short-term OSKM induction reduces the expression of these programs^57^.

Thus, TA →revSC and EC →aVEC shared upregulated adhesion-related programs but differed in their downregulated programs, indicating that the two partially reprogrammed populations arising under the same OSKM induction were accompanied by source-dependent transcriptional changes, as observed for fibroblast source states (Figure 3).

## DISCUSSION

Reprogramming experiments carry a small but experimentally grounded piece of gene-level information that computational regulator-inference methods have not used: the identity of the factors delivered to induce the transition. FateDriver uses these factors as sparse positive labels within a positive–unlabeled-aware ranking framework^22^ and scores every retained gene relative to a specified source-to-target transition. Two design choices follow from this formulation. Because only three or four factors are known positives and the remaining genes are untested, unlabeled genes are ranked rather than treated as negatives, which keeps uncharacterized genes available as candidates. Because the prior network informs gene representations without restricting which genes are scored, FateDriver ranks the full retained transcriptome, whereas network-based methods returned partial rankings of 185–2,453 of the 3,000 evaluated genes.

The benchmark comparisons were designed to exclude specific alternative explanations for this performance. The supervision-matched MLP received the same single-cell data, pseudo-time, and gene-level labels but no graph-based representations, and recovered external reference genes less well in all three datasets; the advantage of FateDriver is therefore not attributable to supervision alone. The MLP also selected the genes most strongly co-expressed with the delivered factors in every dataset, yet recovered fewer reference genes, indicating that proximity to factor expression is not sufficient to identify curated regulators. FateDriver like-wise selected genes with lower median prior-network out-degree than CEFCON and SwitchTFI, which rank genes by network influence or centrality, so its rankings are not explained by network connectivity. Recovery of the protein targets of chemical reprogramming cocktails, which are independent of both the delivered factors and the curated regulator sets, provides a third line of external evidence in the OSKM systems.

Recovery of chemical targets was lower than CEFCON in the cardiac dataset. Two features of that evaluation may contribute. GMT-driven direct conversion and chemical cardiac reprogramming proceed through different cellular routes^9,41^, so the two interventions need not converge on the same genes. In addition, because too few experimentally supported cardiac regulators were present in the analyzed gene space^28^, the cardiac reference set consisted of cardiomyocyte-identity genes, which report the resulting cell state rather than its regulation. The cardiac benchmark therefore tests a different quantity than the OSKM benchmarks, and the two should not be read as equivalent.

The genes prioritized by FateDriver depended on both endpoints of the specified transition. With a common iPSC target, two fibroblast source states shared 18 of their top-100 genes, and the source-specific candidates differed in the direction of their expression change, in inferred regulon activity, and in pathway enrichment; within the source state closer to iPSC, expression of both candidate sets tracked CellRank iPSC fate probability in the directions predicted by their pseudotime trends. With a common fibroblast source, two endpoints shared 38 of their top-100 genes, separating adhesion-associated candidates for the partial-reprogramming endpoint from proliferation-associated candidates for the iPSC-like endpoint. Rankings conditioned on a transition therefore cannot be recovered from the target state alone, which is the practical argument for specifying both populations when prioritizing candidate regulators.

Conditioning on the partial-reprogramming endpoint prioritized *DDX58*, which encodes the double-stranded RNA sensor RIG-I^53^ and was absent from the iPSC-like ranking. At day 10, ISG expression was inversely associated with endogenous pluripotency-marker expression in both donors after adjustment for sequencing depth. In this dataset OSKM was delivered by Sendai virus^25^, and reducing interferon induction has been shown to stabilize Sendai-vector expression^58^. We therefore hypothesize that the endpoint-specific ranking of *DDX58* reflects early antiviral responses that limit sustained OSKM expression, whereas the iPSC-like ranking emphasizes later programs associated with establishing pluripotency^56^. This interpretation is specific to vector-delivered reprogramming and requires perturbation experiments to test.

Extending the same framework to candidate source–target pairs addresses a different question: which population gives rise to a reprogrammed state observed *in vivo*. For two partially reprogrammed intestinal populations arising in the same OSKM-induced tissue, gene-set recovery, residual source identity, nearest-neighbor label transfer, and OSKM-induced developmental potential each ranked TA highest for revSC and EC highest for aVEC. Three of these four read-outs do not involve FateDriver, so the agreement is not internal to the model; it nonetheless rests on transcriptional similarity and plasticity rather than on lineage information, and the two populations are nominated as candidates for lineage tracing rather than established as origins.

Across OSKM-driven pluripotency induction, GMT-driven cardiac conversion, and *in vivo* partial reprogramming, FateDriver converts the factors already delivered in a reprogramming experiment into ranked, transition-specific hypotheses spanning the retained transcriptome. The framework applies to any system in which a source population, a target population, and a set of delivered factors can be defined, and its outputs are intended as prioritized candidates for perturbation experiments.

### Limitations of the study

The predicted regulators and candidate source populations were not experimentally validated. Perturbation experiments are needed to test whether the predicted genes alter reprogramming efficiency or outcome, and genetic lineage tracing is needed to test the source predictions.

Evaluation relied on external gene sets that are incomplete and biased toward well-studied genes, so recovery measures the agreement of a ranking with existing knowledge rather than its accuracy for uncharacterized genes. For GMT-driven conversion, too few experimentally supported regulators were present in the analyzed gene space, and the reference set therefore consisted of cardiomyocyte-identity genes, which report cell identity rather than causal regulation.

FateDriver requires a defined source population, a defined target population, and a set of delivered factors. It is therefore not directly applicable to transitions without experimentally delivered factors, such as development or disease progression, although the same formulation could in principle accept other sets of established regulators as labels. Scoring also depends on an inferred pseudotime ordering, which constrains the cell graph and contributes to the transition prior, so rankings may be sensitive to the trajectory method used.

The contribution of individual model components was not assessed by ablation, and the comparisons reported here evaluate the complete framework against existing methods and a supervision-matched model rather than against reduced versions of itself.

## RESOURCE AVAILABILITY

### Lead contact

Requests for further information and resources should be directed to and will be fulfilled by the lead contact, Haeseung Lee.

### Materials availability

This study did not generate new unique reagents.

## Data and code availability

- This paper analyzes existing, publicly available data. These scRNA-seq datasets are available from the Gene Expression Omnibus under accession numbers GSE181451, GSE23717 GSE242423, GSE297234 and GSE98567; accession numbers are also listed in the key resources table.
- All original code (the FateDriver framework and analysis scripts) is available at https://github.com/Jieun-Sung/FateDriver and is publicly available as of the date of publication. The processed single-cell objects underlying the figures have been deposited at Zenodo (https://doi.org/10.5281/zenodo.21771206). Both are listed in the key resources table.
- Any additional information required to reanalyze the data reported in this paper is available from the lead contact upon request.

## ACKNOWLEDGMENTS

This work was supported by the National Research Foundation of Korea (NRF) grant funded by the Korea government (MSIT) (Nos. RS-2024-00432867, RS-2026-25607354, and RS-2026-45025600097); Glocal University 30 Project at Pusan National University funded by the Ministry of Education(MOE) (2026-glocal-02-004-511-006); and the Ewha Global Excellence Program (EGEP) of Ewha Womans University.

## AUTHOR CONTRIBUTIONS

J.S. and K.S. conceived and designed the study and model architecture and performed down-stream validation analyses. J.S. developed and implemented FateDriver and performed model training and benchmarking. K.S. curated and preprocessed the single-cell datasets. W.K. supervised model development. H.-J.C. provided guidance on reprogramming biology. H.L. supervised dataset construction, model development, and manuscript preparation. J.S. and K.S. wrote the initial draft. W.K., H.-J.C., and H.L. secured funding. All authors reviewed, edited, and approved the final manuscript.

## DECLARATION OF GENERATIVE AI AND AI-ASSISTED TECHNOLOGIES IN THE WRITING PROCESS

During the preparation of this work the authors used Claude and Codex in order to draft and edit sections of the manuscript and to assist in writing analysis code. After using these tools, the authors reviewed and edited the content as needed and take full responsibility for the content of the publication.

## SUPPLEMENTAL INFORMATION INDEX

Figure S1. The reprogramming systems and datasets used in the benchmark, with the embedding and pseudotime ordering of each, related to Figure 2. (A–C) GSE297234 (A), GSE242423 (B) and GSE98567 (C); left, UMAP colored by sample group; right, the same embedding colored by pseudotime.

Figure S2. Overlap and prior-network connectivity of top-100 gene sets, related to Figure 2. (A– C) GSE297234 (GM23815), GSE242423, and GSE98567, respectively. Left, UpSet plots of the six top-100 sets. Bars show the number of genes in each exact intersection, and colored dots indicate the contributing methods. Up to 25 intersections are displayed per dataset. Colors identify the methods throughout. Right, NicheNet out-degree distributions for each method’s selected genes with at least one outgoing interaction. Points represent genes; boxes show medians and interquartile ranges, with whiskers extending to the most extreme values within 1.5 interquartile ranges. The axis is logarithmic, and *n* gives the number of included genes.

Figure S3. Source-state signatures and differential-expression analysis using GSE181451, related to Figure 3. (A) Expression of MEF-enriched genes from Huyghe et al.^10^, HALLMARK EMT, SenMayo senescence, WONG ESC-core, and HALLMARK E2F-target gene sets in MEF-1, MEF-2, and iPSC cells. Violin plots show per-cell mean log-normalized expression after downsampling to 500 UMIs per cell; connected points indicate group means. (B, C) MA plots comparing iPSC with MEF-1 (B) or MEF-2 (C). Each point represents a gene. Orange and blue indicate higher and lower expression in iPSC, respectively, at log_2_ |FC |≥2.5 and adjusted *p <* 0.01; other genes are grey. (D, E) KEGG enrichment of the differentially expressed genes in the corresponding comparisons. Bar lengths indicate− log_10_(FDR); dashed lines mark FDR = 0.01. Bold labels mark pathways discussed in the text.

Figure S4. Clustering and biological programs associated with reprogramming, related to Figure 4. (A) UMAP of GSE297234 cells from donor GM23815, colored by sampling day (D0, D3, D7, and D10). (B) Module scores for six reprogramming-associated programs on the same embedding (red, positive; blue, negative); the gene sets are given in Table S5.

Figure S5. Characterization of the GSE237173 intestine model, related to Figure 5. (A) UMAP of intestinal epithelial cells from Cont and Dox mice, colored by cell type. CBC, crypt base columnar cell; TA, transit-amplifying cell; EC, enterocyte; aVEC, atrophy-induced villus epithelial cell-like cell; revSC, revival stem cell-like cell; EE, enteroendocrine cell; GC, goblet cell; PC, Paneth cell; TC, tuft cell. (B) Cell-type composition in Cont and Dox. (C) Dot plot of marker gene expression for each cell type. Dot size, percentage of expressing cells; color, scaled average expression. (D) Module scores for partial reprogramming and pluripotency in Cont and Dox. *P* values by Wilcoxon rank-sum test. (E) YAP and fetal signature scores on the UMAP embedding, both signatures taken from Kim et al.^54^. (F,G) Venn diagrams of genes upregulated or downregulated in the TA-revSC (F) and EC-aVEC (G) lineages (L1) relative to the alternative lineages (L2 and L3). Genes in the intersection were used for enrichment analysis (Figure 5H, I).

Table S1. Mean rank of the delivered factors for every method and dataset, with gene-level bootstrap confidence intervals, related to Figure 2.

Table S2. Recovery AUROC of every method recomputed after removing curated positives that are also in the annotated-target tier, related to Figure 2.

Table S3. NicheNet outgoing degree of each method’s top-100 genes, related to Figure S2.

Table S4. Shared and source-specific FateDriver candidates in GSE181451, with rank percentiles and expression changes along pseudotime, related to Figure 3.

Table S5. Gene sets for biological-program scoring, day-10 outcome classification, and ISG scoring, related to Figures 4 and 5.

Table S6. Partial- and iPSC-endpoint candidate sets, GO enrichment, donor-specific AUROCs, and ISG-score strata and regression results, related to Figure 4.

Table S7. Candidate-source readouts for the aVEC and revSC populations of GSE237173: gene-set recovery by each candidate’s FateDriver model with bootstrap intervals and paired comparisons, and the expression-based readouts with permutation *p*, related to Figure 5.

Table S8. Public datasets, analysis subsets, and input cell counts used in this study, related to Figures 2–5.

Table S9. Glossary of the symbols used in the STAR Methods, related to Figure 1.

Table S10. Stage-dependent parameters of the training curriculum, related to Figure 1.

Table S11. Co-expression with the delivered-factor module and recovery of external reference gene sets, related to Figure 2.

Table S12. Small molecules of the chemical reprogramming cocktails, their protein targets, and recovery of those targets by every method, related to Figure 2.

## MAIN TABLES, INCLUDING TITLES AND LEGENDS

None.

## SUPPLEMENTAL TABLES

Tables S1–S12 are provided as separate Excel files.

## STAR METHODS

### Key resources table

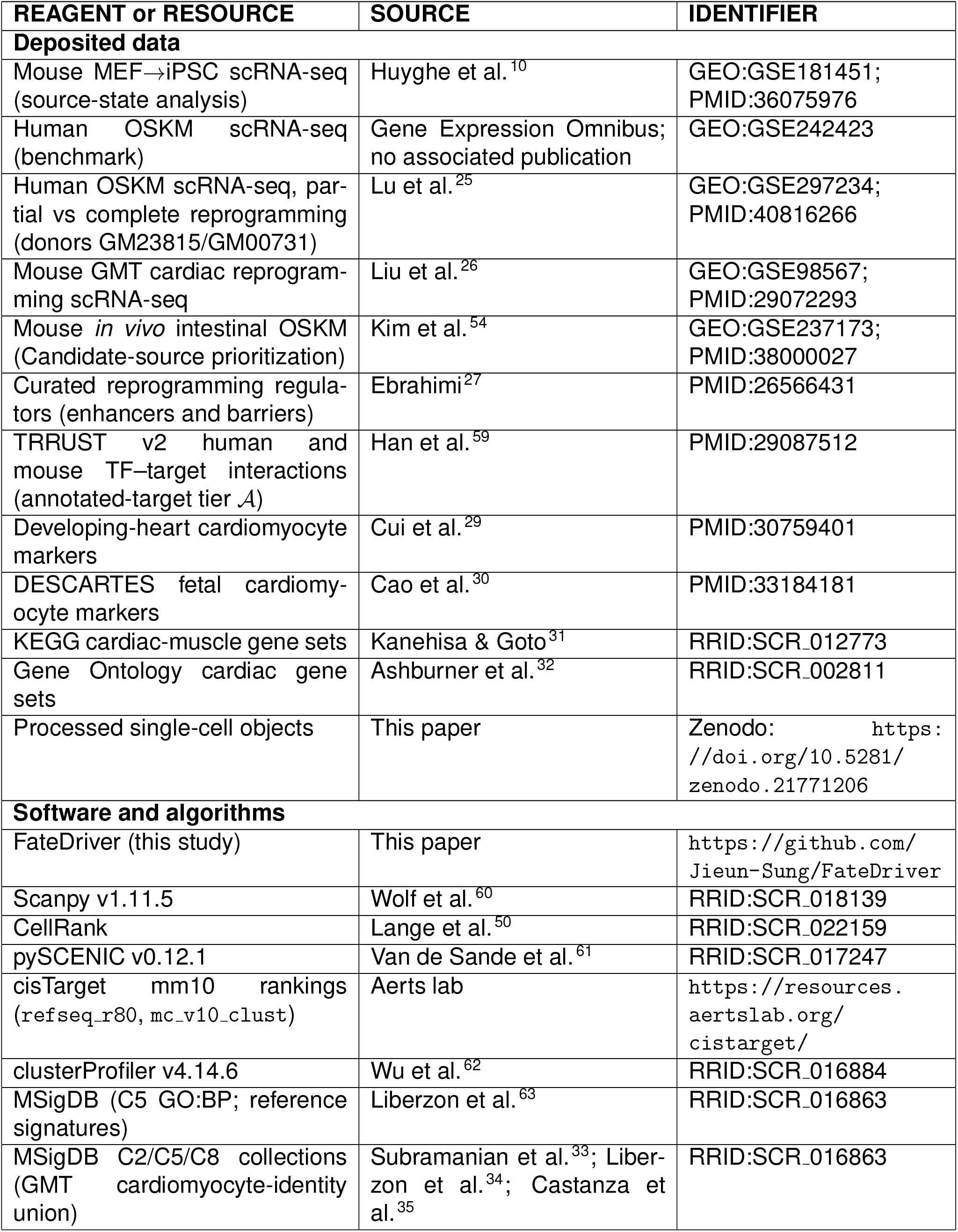
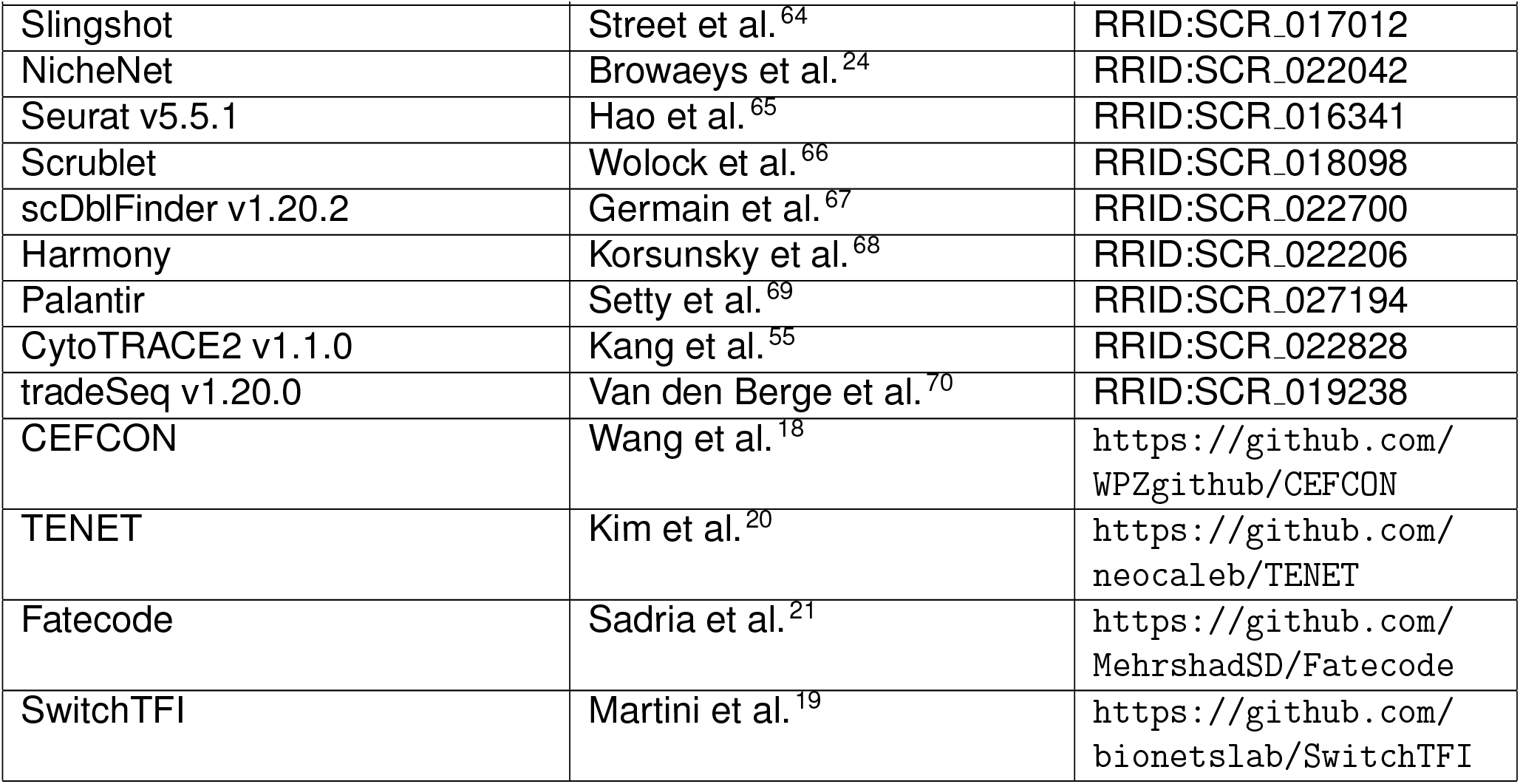

| REAGENT or RESOURCE | SOURCE | IDENTIFIER |
|---|---|---|
| **Deposited data** |  |  |
| Mouse MEF→iPSC scRNA-seq (source-state analysis) | Huyghe et al.<sup>10</sup> | GEO:GSE181451;<br>PMID:36075976 |
| Human OSKM scRNA-seq (benchmark) | Gene Expression Omnibus; no associated publication | GEO:GSE242423 |
| Human OSKM scRNA-seq, par-tial vs complete reprogramming (donors GM23815/GM00731) | Lu et al.<sup>25</sup> | GEO:GSE297234;<br>PMID:40816266 |
| Mouse GMT cardiac reprogram-ming scRNA-seq | Liu et al.<sup>26</sup> | GEO:GSE98567;<br>PMID:29072293 |
| Mouse *in vivo* intestinal OSKM (Candidate-source prioritization) | Kim et al.<sup>54</sup> | GEO:GSE237173;<br>PMID:38000027 |
| Curated reprogramming regula-tors (enhancers and barriers) | Ebrahimi<sup>27</sup> | PMID:26566431 |
| TRRUST v2 human and mouse TF–target interactions (annotated-target tier 𝒜) | Han et al.<sup>59</sup> | PMID:29087512 |
| Developing-heart cardiomyocyte markers | Cui et al.<sup>29</sup> | PMID:30759401 |
| DESCARTES fetal cardiomy-ocyte markers | Cao et al.<sup>30</sup> | PMID:33184181 |
| KEGG cardiac-muscle gene sets | Kanehisa & Goto<sup>31</sup> | RRID:SCR_012773 |
| Gene Ontology cardiac gene sets | Ashburner et al.<sup>32</sup> | RRID:SCR_002811 |
| Processed single-cell objects | This paper | Zenodo: https://doi.org/10.5281/zenodo.21771206 |
| **Software and algorithms** |  |  |
| FateDriver (this study) | This paper | https://github.com/Jieun-Sung/FateDriver |
| Scanpy v1.11.5 | Wolf et al.<sup>60</sup> | RRID:SCR_018139 |
| CellRank | Lange et al.<sup>50</sup> | RRID:SCR_022159 |
| pySCENIC v0.12.1 | Van de Sande et al.<sup>61</sup> | RRID:SCR_017247 |
| cisTarget mm10 rankings (refseq_r80, mc_v10_clust) | Aerts lab | https://resources.aertslab.org/cistarget/ |
| clusterProfiler v4.14.6 | Wu et al.<sup>62</sup> | RRID:SCR_016884 |
| MSigDB (C5 GO:BP; reference signatures) | Liberzon et al.<sup>63</sup> | RRID:SCR_016863 |
| MSigDB C2/C5/C8 collections (GMT cardiomyocyte-identity union) | Subramanian et al.<sup>33</sup>; Liber-zon et al.<sup>34</sup>; Castanza et al.<sup>35</sup> | RRID:SCR_016863 |
| Slingshot | Street et al.<sup>64</sup> | RRID:SCR_017012 |
| NicheNet | Browaeys et al.<sup>24</sup> | RRID:SCR_022042 |
| Seurat v5.5.1 | Hao et al.<sup>65</sup> | RRID:SCR_016341 |
| Scrublet | Wolock et al.<sup>66</sup> | RRID:SCR_018098 |
| scDblFinder v1.20.2 | Germain et al.<sup>67</sup> | RRID:SCR_022700 |
| Harmony | Korsunsky et al.<sup>68</sup> | RRID:SCR_022206 |
| Palantir | Setty et al.<sup>69</sup> | RRID:SCR_027194 |
| CytoTRACE2 v1.1.0 | Kang et al.<sup>55</sup> | RRID:SCR_022828 |
| tradeSeq v1.20.0 | Van den Berge et al.<sup>70</sup> | RRID:SCR_019238 |
| CEFCON | Wang et al.<sup>18</sup> | https://github.com/WPZgithub/CEFCON |
| TENET | Kim et al.<sup>20</sup> | https://github.com/neocaleb/TENET |
| Fatecode | Sadria et al.<sup>21</sup> | https://github.com/MehrshadSD/Fatecode |
| SwitchTFI | Martini et al.<sup>19</sup> | https://github.com/bionetslab/SwitchTFI |

### Experimental model and study participant details

#### Datasets

All analyses used previously published scRNA-seq datasets of cellular reprogramming obtained from the Gene Expression Omnibus; no new experiments were performed. The regulator-recovery benchmark used two human OSKM-induced pluripotency datasets, GSE297234 GSE242423 (no associated publication), and one mouse GMT-induced cardiac reprogramming dataset, GSE98567^26^ (*Gata4, Mef2c, Tbx5*). The endpoint-conditioning analysis used GSE297234 comparing partial-reprogramming and iPSC-like endpoints in a young donor (GM23815, 22 years), with an aged donor (GM00731, 96 years) analyzed for cross-donor reproducibility. The source-state analysis used a mouse MEF →iPSC dataset, GSE181451^10^, in which the starting fibrob-lasts resolved into two transcriptionally distinct states, MEF-1 (*n* = 5,243 cells) and MEF-2 (*n* = 1,505 cells), with iPSC as the common target (*n* = 3,876 cells). The candidate-source analysis used a mouse intestinal dataset with doxycycline-inducible OSKM expression *in vivo*, GSE237173^54^. Preprocessing, annotation, and pseudotime inference are described under STAR Methods, *Datasets and preprocessing*. Sex and other donor covariates were not modeled, because each reprogramming system is defined by its source and target cell identities.

### Method details

#### Datasets and preprocessing

Five publicly available scRNA-seq datasets capturing cellular reprogramming were obtained from the Gene Expression Omnibus (GSE297234, GSE242423, GSE98567, GSE181451, and GSE237173). Three datasets model reprogramming from a fibroblast source toward induced pluripotent stem cells via OSKM—GSE297234 and GSE242423 (human) and GSE181451 (mouse while GSE98567 (mouse) models GMT-induced reprogramming toward cardiomyocytes. GSE2371 (mouse) models doxycycline-inducible OSKM reprogramming of intestinal epithelium and is described separately below. Dataset composition and post-QC cell numbers are summarized in Table S8. Raw count matrices were processed in Scanpy v1.11.5^60^ (GSE297234, GSE98567, GSE242423, and GSE181451) or Seurat v5.5.1^65^ (GSE237173). Genes detected in fewer than 3 cells and cells with fewer than 200 detected genes were removed. Cells were then filtered on detected gene count, total UMI count, and mitochondrial percentage using dataset-specific thresholds from the empirical per-sample distributions (typically 200–7,500 genes, ≥500 UMIs, and *<*20–25% mitochondrial reads). The plate-based GSE98567, with much higher per-cell depth, used correspondingly higher bounds (500–12,000 genes, ≥100,000 UMIs). Doublets were removed with Scrublet (Python)^66^ or scDblFinder v1.20.2 (R)^67^ prior to downstream analysis. Counts were log-normalized, the top 3,000 highly variable genes were selected per dataset, and expression was scaled with regression of mitochondrial percentage across all datasets before principal component analysis. A shared-nearest-neighbor graph was built from the first 30 principal components, with Harmony integration^68^ applied to GSE98567 to correct batch effects across its multiple samples. Leiden clustering was performed with default parameters, though downstream analyses relied on cell-type or sample labels. UMAP embeddings were computed on the same graph for visualization. GSE242423, GSE98567, and GSE181451 were grouped by condition or time point, and GSE297234 was annotated with the reprogramming-state signatures of its original study^25^ (STAR Methods, *Identification of partial reprogramming and pluripotency states*). GSE237173 was annotated with canonical markers from its original study^54^ and analyzed as described under STAR Methods, *Single-cell characterization of OSKM-induced intestinal partial reprogramming*.

Pseudotime was estimated to establish the cellular ordering along the reprogramming process across the fibroblast-source datasets. For GSE242423, GSE98567, and GSE181451, per-cell pseudotime was inferred with Slingshot v2.10.0^64^ on the UMAP embedding, using annotated cell types as cluster labels. The fibroblast cluster was specified as the starting cluster, so that cells within the source population were also ordered along the trajectory, and the re-programmed target cluster (iPSC or cardiomyocyte) was specified as the end cluster. Lineage trajectories were additionally inferred for GSE237173, rooted at its intestinal epithelial source. For GSE297234, whose four time points branch toward partial-reprogramming and iPSC-like states, we used Palantir v1.4.4^69^ to infer pseudotime and branch probabilities, starting from a cell in the day-0 (D0) population.

### The FateDriver model

#### Architecture overview

FateDriver jointly trains two graph encoders, an expression decoder, and a gene scorer (Figure 1A–E). The encoders map cells and genes into a shared space. The decoder reconstructs expression counts from these embeddings, while the scorer relates genes to the source and target populations. Training combines reconstruction, ranking, and shift-consistency losses.

#### Notation

Bold uppercase denotes matrices, bold lowercase vectors, and calligraphic upper-case index sets. The indices *i, g*, and *ϵ* denote cells, genes, and training epochs, respectively. Table S9 provides the symbol glossary.

The inputs are library-normalized, log-transformed expression 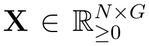, raw counts 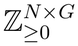, and cell pseudotime values *t*_*i*_. Dataset annotations define source and target populations *S* and *T*. The retained genes comprise the precomputed highly variable set and all delivered factors. Cells and genes share an *H*-dimensional space, with *H* = 1024:

**Z** ∈ ℝ^*N ×H*^ cell embeddings; row **z**_*i*_ is one cell

**E** ∈ ℝ^*G×H*^ gene embeddings; row **e**_*g*_ is one gene

**z**_*S*_, **z**_*T*_ ∈ ℝ^*H*^ the source and target populations pooled into one vector each

**Δ** = **z**_*T*_ − **z**_*S*_ the source-to-target transition used in gene scoring

*s*_*g*_ ∈ ℝ the score of gene *g*, whose ranking is the output of the model

Genes are grouped into delivered factors *D*, their annotated targets *A* (the intermediate tier), and unlabeled genes *U* = {1, … , *G*} \ (*D* ∪ *A* ). Delivered factors are *POU5F1, SOX2, KLF4*, and *MYC* for OSKM reprogramming, and *Gata4, Mef2c*, and *Tbx5* for GMT conversion; they are excluded from *A*. Because unlabeled genes may include uncharacterized regulators, they are not treated as negatives, and FateDriver supervises relative rankings within a positive–unlabeled framework^23^.

#### Cell splitting and evaluation

All FateDriver analyses used cell-based splits. Within each run, source and target cells were shuffled separately using the run seed and divided into disjoint training, validation, and test pools. In the three benchmark datasets, validation and test pools each contained ⌊0.1*n*⌋ cells from a population of size *n*; the remaining cells formed the training pool. The candidate-source analysis used 25% for each evaluation pool. Training episodes sampled training pools. Validation episodes supported model selection and pseudo-positive bank updates, and final scores were averaged over test episodes. Delivered-factor labels were shared across the cell splits. Recovery of external reference gene sets was evaluated separately, as described under *Recovery of external reference gene sets*.

#### Cell features and the trajectory graph

Cell features **C**∈ ℝ;^*N ×*50^ are the top 50 truncated-SVD components of **X**, standardized to zero mean and unit variance^71^. The SVD basis and standardization parameters are fitted on training source and target cells and applied to all cells. Each cell is connected to its 15 nearest neighbors under squared Euclidean distance in **C**. Edges are retained only from earlier to later cells in pseudotime:

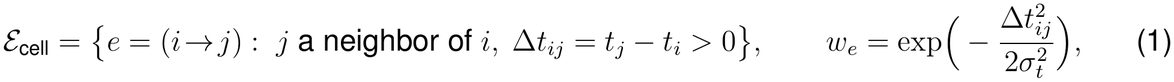

where *σ*_*t*_ is the median |Δ*t*_*ij*_| over retained edges. Weights for duplicate cell-graph edges are summed. The graph includes all cells, including those outside the source and target populations.

#### Cell encoder

Both encoders use two-layer graph convolutional networks (GCNs)^72^ with row normalization for directed, weighted edges. Let 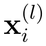 denote the features of node *i* at layer *l*, **W**^(*l*)^ the learned transformation, and *δ*_*i*_ the weighted in-degree. Each layer computes

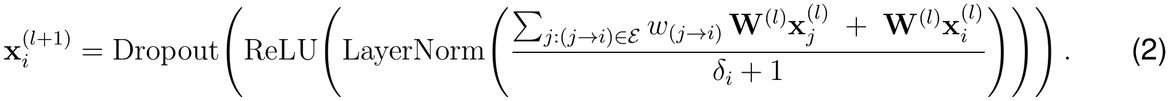

The cell encoder returns **Z** = Enc_cell_(**C**; ε_cell_) with hidden width *H* and dropout rate 0.1. Aggregation combines each cell’s features with those of its incoming neighbors, which precede it in pseudotime.

#### Gene encoder

Gene features are initialized as **X**^*⊤*^**Z** + **E**^(0)^, where **E**^(0)^ ∈ℝ^*G×H*^ contains learnable gene-specific vectors. Layer normalization and dropout precede refinement by the gene GCN. Thus, each gene combines an expression-weighted sum of cell embeddings with a learnable residual. The prior graph combines the NicheNet gene regulatory and signaling networks^24^; ligand– receptor and ligand–target networks are excluded. The human input union contains 33,354 genes and 8,419,668 edges. Edges are restricted to retained genes and kept if at least one endpoint belongs to the 3,000 genes with the highest mean expression. The filtered GSE242423 graph contains 3,000 genes and 191,010 edges. Edge weights are floored at 10^*−*6^ and divided by their maximum. Edge directions are reversed so that genes receive information from their regulatory or signaling targets. Two layers of Equation (2) produce **E**. Genes without prior-network edges retain their expression-derived and learnable features through the self-node term.

#### Edge pruning

After a 20-epoch warm-up, edges are scored every 20 epochs. For an *L*^2^-normalized endpoint embedding 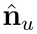 and current mean graph weight 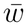,

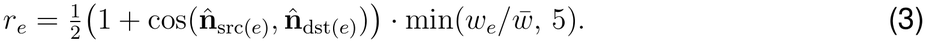

Scores are smoothed across pruning events by an exponential moving average with decay 0.8. Each event removes up to 10% of current cell edges or 5% of gene edges, in ascending score order. Removal is prevented if a node would fall below degree 5 in the cell graph or 3 in the gene graph.

#### Expression decoder and reconstruction loss

The decoder predicts the count-model mean *µ*_*ig*_ from cell and gene embeddings. With a learned bilinear transformation **W**_*B*_ ∈ℝ^*H×H*^ , gene bias *b*_*g*_, and depth coefficient *ν*,

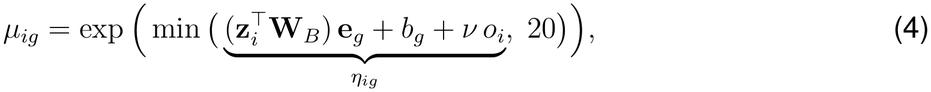

where *o*_*i*_ = log max(Σ_*g*_ *K*_*ig*_, 1) is the cell’s total-count offset. Raw counts are modeled by a zero-inflated negative binomial (ZINB) distribution^73^, with gene-specific dispersion *θ*_*g*_ and structural-zero probability *ζ*_*g*_. The matrix **W**_*B*_ is initialized to the identity, and *b*_*g*_ to mean log-normalized expression. Dispersion *θ*_*g*_ and structural-zero probability *ζ*_*g*_ are softplus and sigmoid transformations of unconstrained gene-specific parameters, initialized near 2 and 0.1, respectively. For the negative binomial distribution NB, the log-likelihood is

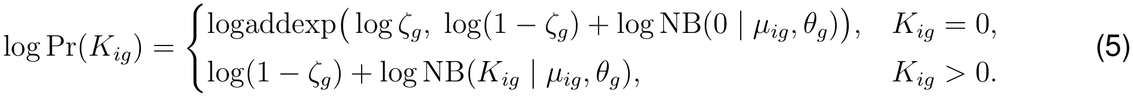

The reconstruction loss is

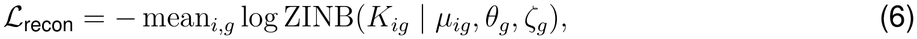

where the mean spans min(*N*_train_, 5000) sampled training cells and all *G* genes. This loss updates both encoders; the decoder output does not enter the gene score.

#### Population encoding and the transition vector

Each episode samples 128 source and 128 target cells, with replacement when a population is smaller. A shared, permutation-invariant attention pooling module^74^ summarizes each sample.

For sampled cell embeddings **Z**_*\**_, the set encoder uses a learnable seed **q** ∈ℝ^*H*^ , *n*_*h*_ = 4 attention heads, and three feed-forward blocks FF_1,2,3_. With multi-head attention MHA and layer normalization LN,

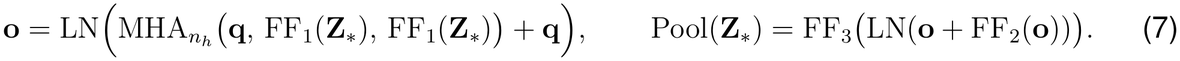

Applying this module separately to source and target samples gives **z**_*S*_ and **z**_*T*_ . Their difference defines the transition:

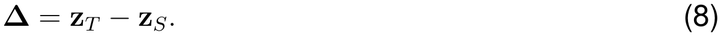

#### Gene scoring head

Before scoring, multi-head self-attention^75^ produces contextualized gene embeddings 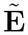. Each gene attends to the genes in its 512-gene chunk and a global mean gene token. A residual connection and layer normalization yield the updated embedding 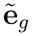. The scorer combines cross-attention, similarity, and prior terms:

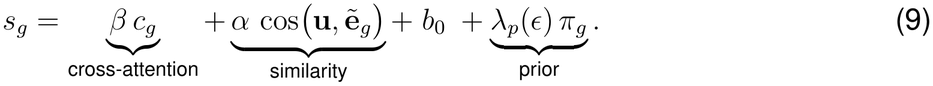

Here *α, β*, and *b*_0_ are learned scalars, and *λ*_*p*_(*ϵ*) is the scheduled prior weight.

*Cross-attention term*. The source, target, and transition form three tokens, **M** = LN([**z**_*S*_; **z**_*T*_ ; **Δ**])ℝ^3*×H*^ , where LN denotes layer normalization. Linear projections map each gene to a query and each token to a key and value. For head *h* among *n*_*h*_ = 4 heads, these vectors are **q**_*gh*_, **k**_*ah*_, and 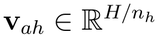 , with *a* ∈ {*S, T*, Δ}. A learned read-out vector ***κ***_*h*_ maps each value to a scalar:

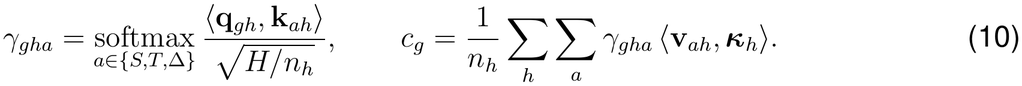

The gene-specific attention weights combine the three token readouts, and *c*_*g*_ averages these contributions across heads.

*Similarity term*. An MLP maps the concatenated source, target, and transition vectors to a normalized control vector:

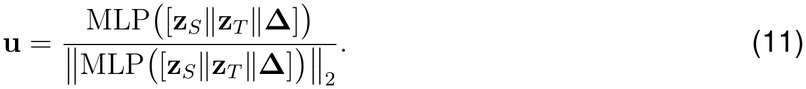

The term cos 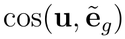 measures gene alignment with this direction.

*Prior term*. Five gene-level features supplement the embedding-based terms: positive target-minus-source differences in mean expression and detection fraction, absolute mean-expression difference, gene-graph out-degree, and positive correlation with pseudotime. Each feature is mapped to [0, 1] by rk(*x*)_*g*_ = (rank(*x*_*g*_) − 1)*/*(*G*− 1), forming ***ϕ***_*g*_ ∈ℝ^5^. Softmax-constrained learned weights ***ω*** and bias *b*_*π*_ give

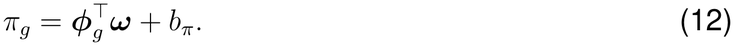

The exported score is the mean of *s*_*g*_ over ten resampled episodes drawn from test cells.

#### Three-tier ranking objective

The objective supervises *D* ≻ *A* ≻ *U*, assigning delivered factors higher priority than their downstream targets to distinguish the experimental perturbation from its response. The annotated-target tier (intermediate tier) contains targets of the delivered factors annotated in the species-matched TRRUST v2 table^59^. For higher-priority set *P*, lower-priority set *N*, and margin *m*, the pairwise loss is

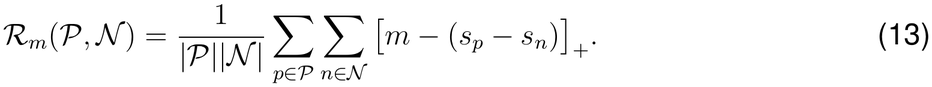

Each step samples 500 comparison genes. A fraction *f*_*↓*_(*ϵ*) comes from *A* and the pseudo-positive bank *B*, forming *N*_*↓*_. The remainder, *N*_pure_, is sampled uniformly from *U* \ *B*. The loss combines the three comparisons:

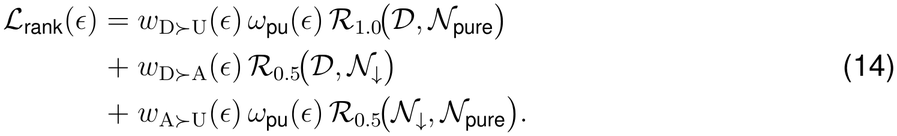

The PU factor reduces penalties involving unlabeled genes:

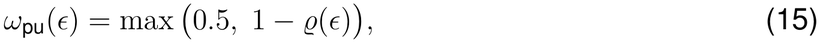

where *ϱ* increases from 0 to 0.30, so *ω*_pu_ decreases from 1 to 0.70. The floor at 0.5 is not reached. Tier weights and sampling fractions follow Table S10. From epoch 10, a pseudo-positive bank is updated using validation-averaged gene scores 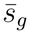. Bank members join the intermediate tier and are excluded from the unlabeled sampling pool. Similarity to labeled genes is

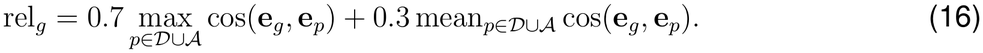

At each update, an unlabeled gene accrues a hit when its score and relation value meet or exceed the 99.5th and 98th percentiles, respectively, over *U*. The hit count *h*_*g*_ increases by one for qualifying genes and is otherwise unchanged. The promotion statistic is updated as

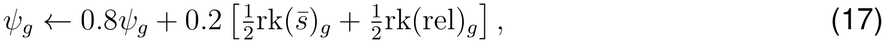

where each rank transform is computed across the scored genes. Among unlabeled genes with *h*_*g*_≥ 2, up to 128 are selected in descending order of *ψ*_*g*_ + 0.02*h*_*g*_. The bank is rebuilt at every validation pass. Eligibility therefore depends on cumulative hits rather than consecutive qualifying epochs. Genes leaving the bank become eligible for unlabeled sampling again.

#### Shift-consistency loss

The shift-consistency loss aligns the control vector, the mean embedding of top-ranked genes, and individual top-ranked gene embeddings with **Δ**. Let *K* contain the 128 highest-scoring genes and **ê**_*g*_ denote normalized gene embeddings before self-attention. Score weights at temperature 0.2 and their normalized mean embedding are

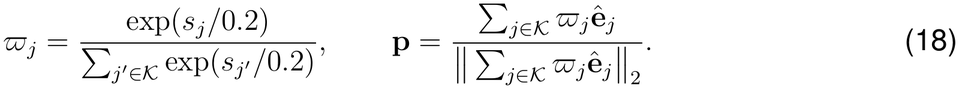

With 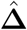 denoting the normalized transition, the loss is

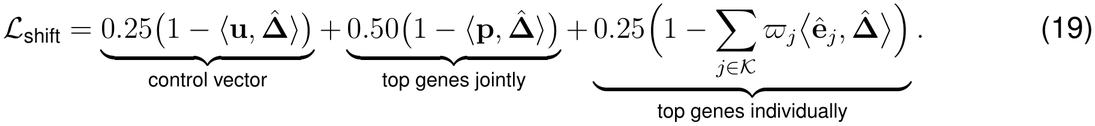

#### Total loss and training schedule

The total loss is

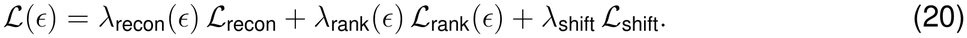

All three losses remain active throughout training. Over a maximum of *E* = 200 epochs, the weights follow a cosine schedule, with 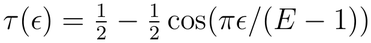:

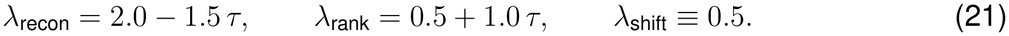

The reconstruction weight decreases from 2.0 to 0.5, the ranking weight increases from 0.5 to 1.5, and the shift-consistency weight remains 0.5.

The ranking curriculum has three stages. Stage 1 (epochs 0–49) supervises only *D* ≻ *U* . Stage 2 (epochs 50–69) introduces the annotated-target tier and increases PU relaxation *ϱ* from 0 to 0.09. Stage 3 (epochs 70–199) increases *ϱ* from 0.09 to 0.30. Within stages 2 and 3, scheduled quantities interpolate linearly between their first- and last-epoch values. These include prior scale *λ*_*p*_, annotated-target sampling fraction *f*_*↓*_, tier weights *w*_D*≻*A_ and *w*_A*≻*U_, and *ϱ* (Table S10). Margins remain 1.0 for delivered-versus-unlabeled comparisons and 0.5 for adjacent tiers.

Optimization uses Adam with learning rate 10^*−*4^ and weight decay 10^*−*4^ under automatic mixed precision. Each epoch has 10 steps of 1,024 episodes. From stage 3 onward, validation AUPRC over *D* ∪ *A* determines model selection. Training stops after 10 epochs without improvement, and the best checkpoint is restored. Early stopping can truncate the cosine schedule before its final epoch.

#### Benchmark methods

We compared FateDriver with a supervision-matched MLP and four published methods: CEF-CON, TENET, Fatecode, and SwitchTFI. The published methods do not use delivered factors as gene-level labels and were run with the parameters recommended by their authors unless stated otherwise. All rankings were restricted to a common evaluation set of 3,000 highly variable genes per dataset, and each method was evaluated on the genes it scored. Source and target populations were D0 and iPSC cells in GSE242423, D0 and D10 cells of donor GM23815 in GSE297234, and uninfected control and GMT-reprogrammed cells in GSE98567.

#### Supervision-matched MLP

Each episode sampled 128 source and 128 target cells. For each gene, the input comprised its expression in these cells and their pseudotime values (512 features), with pseudotime features shared across genes within an episode. Standardization was fitted to training-episode feature rows, and a feed-forward network with hidden layers of 256 and 64 units mapped each vector to a scalar score. The MLP used the same cell splits, delivered-factor labels, and annotated-target labels as FateDriver. Its pairwise loss enforced delivered factors ≻ annotated targets ≻remaining genes, without positive–unlabeled relaxation. Annotated targets were excluded from internal validation and test metrics, and external recovery followed the common evaluation procedure below.

#### CEFCON^18^

CEFCON constructs a lineage-specific GRN from expression and a prior network, then applies network control theory. We ran CEFCON for each dataset-specific source–target comparison with min–max-normalized pseudotime. The prior was the human NicheNet interaction network for GSE242423 and GSE297234 and its mouse-converted network for GSE98567. GRN learning used 3,000 highly variable genes and 350 training epochs. Outputs were aggregated over three independent repeats, and edges were thresholded at an average degree of 8. The per-gene network-influence score determined gene rankings.

#### TENET^20^

Source and target cells were ordered by pseudotime for transfer-entropy-based GRN inference. Transfer entropy was estimated for each ordered gene pair with history length 1 and converted to *z*-scores and *p* values. Edges were filtered at Benjamini–Hochberg FDR *<* 0.01, followed by removal of indirect edges. Network out-degree determined gene scores.

#### Fatecode^21^

The classifier-guided autoencoder was trained on source and target cells using cell-type annotations as fate labels. Each latent dimension was scanned over a grid of scaling factors. We selected the perturbation that maximally increased the predicted target population. Gene scores were obtained by decoding the difference between the selected perturbed and original latent codes and extracting the target-fate component.

#### SwitchTFI^19^

SwitchTFI uses regression stumps to evaluate TF–target interactions against cell-state labels. We supplied binary source/target labels for each dataset without pseudotime. The prior GRN was the NicheNet human network for GSE242423 and GSE297234 and the mouse network for GSE98567. Edges were pruned for insufficient co-expression and retained without additional family-wise-error filtering. PageRank centrality in the transition GRN determined TF rankings.

### Benchmarking and downstream validation analyses

#### Consensus top-100 gene selection

FateDriver and the MLP were each trained with five random seeds (42–46). Within each run, rank *r* among *n* genes was converted to 1− (*r*− 1)*/*(*n*− 1), with higher values indicating higher ranks. Genes were ordered by their mean percentile across runs, with ties resolved by gene symbol. Each published method contributed one final ranking. Rankings were re-ranked within each method’s scored subset of the evaluation set. Overlap and network-connectivity analyses used the first 100 genes, retaining delivered factors. Co-expression used the first 100 genes after removing delivered factors; recovery used the full rankings after this removal.

#### Ranking of the delivered factors

We calculated the mean rank of OSKM in GSE297234 and GSE242423, and of GMT in GSE98567. Missing factors were assigned rank 3,000. Confidence intervals were obtained by bootstrapping factor genes, using the common procedure under *Quantification and statistical analysis* (Table S1).

#### Curation of external reference gene sets

For OSKM datasets, the curated reprogramming regulators comprised 17 enhancers and 35 barriers from Ebrahimi^27^. Included genes increased iPSC efficiency or kinetics upon activation or overexpression (enhancers), or upon depletion or inhibition (barriers). Aliases were resolved as p53 to *TP53*, p21 to *CDKN1A*, p16Ink4a/p19Arf to *CDKN2A*, and C/EBP*α* to *CEBPA*. Non-gene entries, including vitamin C, were excluded.

For cardiac reprogramming, we used cardiomyocyte-identity signatures from the Cui and DESCARTES atlases^29,30^, together with KEGG and Gene Ontology sets^31,32^. These comprised KEGG *cardiac muscle contraction* and GO *cardiocyte differentiation, cardiac muscle cell differentiation, striated muscle contraction*, and *sarcomere organization*. Identity sets were obtained from MSigDB^33–35^. OSKM regulator recovery and cardiac identity-gene recovery were evaluated separately.

#### Recovery of external reference gene sets

The area under the receiver operating characteristic curve (AUROC) measured recovery of reference genes from each method’s scores. Delivered factors were excluded, and the evaluated universe comprised the method’s scored genes within the common evaluation set. Confidence intervals used stratified bootstrap resampling of reference and non-reference genes separately (Table S11). To assess overlap with training labels, we repeated each method’s analysis after removing annotated targets (TRRUST) from both the reference set and the evaluated rankings (Table S2).

#### Chemical-cocktail target sets

Protein targets were collected for compounds used in five pluripotency protocols^36–40^ and six cardiac protocols^41–46^. Compounds were matched to Drug-Bank by accession, PubChem identifier, or normalized name or synonym^76^. DrugBank accessions in PubChem synonym lists supplemented incomplete cross-references. Human targets were retrieved from DrugBank^77^ and BindingDB^78^. For each BindingDB record, we selected the minimum available *K*_i_, *K*_d_, or IC_50_, then retained compound–protein pairs with a median affinity of at most 1 *µ*M across records. UniProt accessions were mapped to primary gene symbols^79^, and symbols were uppercased for matching, including in the mouse cardiac dataset.

The main analysis used the union of DrugBank therapeutic targets and affinity-filtered BindingDB targets. Sensitivity analyses used DrugBank targets alone or BindingDB cutoffs of 0.1 and 10 *µ*M. Recovery followed the common AUROC procedure. Overlapping TRRUST annotated-target labels were retained in the main analysis and removed in a separate analysis. The pluripotency and cardiac collections contained 33 and 23 compounds, respectively. Therapeutic-only target sets contained 117 and 90 genes; unions with affinity-filtered BindingDB targets contained 262 and 130. Compound identifiers and target lists are provided in Table S12. Overlap with TRRUST annotated labels is reported as gene counts only.

#### Co-expression with the delivered-factor module

The delivered-factor module was scored per cell from normalized expression using `scanpy.tl.score genes`. Cells with finite pseudotime were partitioned into 20 equal-count bins, or fewer if quantile boundaries coincided. For each selected gene, we calculated the Pearson correlation between its bin-mean expression and the bin-mean module score. The method-level measure was the mean correlation across its top-100 genes, selected after removing delivered factors. Confidence intervals used gene-level bootstrap resampling of these correlations (Table S11).

#### Overlap and prior-network connectivity of prioritized genes

Exact intersections were calculated among the six methods’ top-100 sets, retaining delivered factors. The largest 25 intersections were displayed in each UpSet plot. Out-degree was the number of outgoing interactions in the NicheNet regulatory-and-signaling network. Degree statistics included only genes with at least one outgoing interaction. We calculated the median, interquartile range, and a gene-level bootstrap interval for the median (Table S3).

#### Source-state signature analysis in GSE181451

We compared MEF-enriched, EMT, senescence, ESC-core, and E2F-target signatures across MEF-1, MEF-2, and iPSC cells. The MEF signature comprised the 150 genes for clusters 1 and 2 in Supplementary Table 1 of the original dataset publication^10^. The remaining signatures came from MSigDB^33,63^: HALLMARK EPITHELIAL MESENCHYMAL TRANSITION, SAUL SEN MAYO, WONG EMBRYONIC STEM CELL CORE, and HALLMARK E2F TARGETS. Human symbols were converted to mouse-style capitalization, and duplicates were removed.

Raw counts were downsampled to 500 UMIs per cell without replacement by multivariate hypergeometric sampling. All cells in the three populations were retained. Counts were normalized to 10,000 per cell and transformed by log(1 + *x*). Each cell’s signature score was the mean transformed expression across matched genes.

#### Direction of expression change along pseudotime

Source-specific candidates were profiled along pseudotime, excluding genes shared by the two top-100 sets. Cells were divided into 50 equal-occupancy bins. For each gene, we calculated mean log-normalized expression within each bin. We smoothed these values by averaging each bin with the two preceding and two following bins. At the boundaries, values were extended by reflection to retain a five-bin window. Each gene’s smoothed profile was min–max scaled to 0–1 across all pseudotime bins. Mean relative expression and its standard error were calculated across all genes in each source-specific set at each bin.

#### Regulon activity

Regulon activity was computed with pySCENIC v0.12.1^61^ using GSE181451 counts and genes detected in at least 1% of cells. GRNBoost2 inferred co-expression modules from mouse TFs. Motif pruning used mm10 cisTarget rankings `refseq r80` and `mc v10 clust`. The ranking databases covered 500 bp upstream to 100 bp downstream of transcription start sites and 10 kbp on either side. Pruning used `motifs-v10nr clust-nr.mgi` annotations with dropout masking. Per-cell AUCell scores were averaged across motif-supported regulons in each candidate set. Control MEF and OSKM-induced iPSC distributions were compared using two-sided Kolmogorov–Smirnov tests.

#### Pathway analysis

MEF-1-specific and MEF-2-specific genes were tested separately for KEGG over-representation with clusterProfiler v4.14.6^62^ (`enrichKEGG`, organism `mmu`). Symbols were mapped to Entrez identifiers using `org.Mm.eg.db`. The background comprised KEGG-annotated mouse genes, with pathway sizes restricted to 10–500. *P* values were adjusted by the Benjamini– Hochberg procedure. Human Diseases pathways were excluded, and up to ten pathways per candidate set were displayed at FDR *<* 0.05. Enrichment factor was the pathway fraction in the query divided by that in the background.

Pathways were displayed in decreasing −log_10_(FDR), with ties resolved by enrichment factor. Differential-expression pathway plots used the same FDR ordering.

#### Differential expression between iPSC and each source state

We used the iPSC-versus-MEF-1 and iPSC-versus-MEF-2 differential-expression comparisons, with positive log_2_ fold changeindicating higher expression in iPSC. Genes with |log_2_ FC |≥2.5 and adjusted *p <* 0.01 were selected for KEGG over-representation analysis.

#### Fate-probability analysis using CellRank

CellRank^50^ was applied to cells with finite Sling-shot pseudotime using the shared neighbor graph with 15 neighbors and 30 principal components. The transition matrix combined pseudotime and connectivity kernels with weights 0.8 and 0.2. GPCCA used 20 Schur vectors and six macrostates, with cell-state annotations as the cluster key. Terminal states were predicted automatically. The terminal macrostate with the highest fraction of iPSC-annotated cells defined the iPSC endpoint. Absorption probabilities were calculated once and reused across comparisons.

Source-state probabilities were compared with a one-sided Mann–Whitney *U* test for higher values in MEF-2. Bimodality was assessed using the bimodality coefficient (threshold 0.555) and Hartigan’s dip test. MEF-2 cells with iPSC absorption probability ≥ 0.5 formed the iPSC-high fraction; the remainder formed the iPSC-low fraction.

MEF-1-specific and MEF-2-specific sets were scored with `scanpy.tl.score genes`, excluding OSKM factors. Each score was the mean log-normalized expression of candidate genes minus that of expression-matched control genes, using the AnnData `raw` slot. The control-set size was the greater of 50 and the number of matched candidate genes (79 and 75 for the two directional sets). Module scores were compared between the two MEF-2 fractions using two-sided Mann– Whitney *U* tests.

#### Identification of partial reprogramming and pluripotency states

Cells were clustered with Leiden in Scanpy (resolution 0.3). Reprogramming-state signatures from Lu et al.^25^ were scored with `scanpy.tl.score genes`. Each cluster was assigned to the state with the highest mean module score, defining the fibroblast source and partial-reprogramming and iPSC endpoints. Additional MET, EMT, epigenetic-reset, cell-cycle, DNA damage response (DDR), and senescence signatures were scored by the same procedure (Table S5).

#### Endpoint-specific candidate selection

FateDriver was trained separately for fibroblast →partialand fibroblast→ iPSC transitions in GM23815 using five seeds (42–46). Genes scored in all runs were retained. Within each run, ranks were divided by the maximum rank. The displayed percentile was one minus the mean of these values across runs. Rank-percentile heatmaps used a nonlinear color scale to emphasize upper ranks. The top 100 genes for each endpoint were selected from the 1,767 genes common to both rankings. Their intersection defined the shared set; genes selected for only one endpoint defined endpoint-specific sets. Overlap was tested by an upper-tail hypergeometric test with the common genes as background (Table S6).

#### GO enrichment of endpoint-specific candidates

Shared and endpoint-specific sets were tested against MSigDB C5 GO biological processes (human symbols, v2025.1)^63^. GO sets were restricted to the 1,767 common genes, retaining sets of 10–500 genes. Upper-tail hypergeometric *p* values were adjusted within each candidate set by the Benjamini–Hochberg procedure. Enrichment factor was (*k* + 1)*/*(*nK/N* + 1), where *k* is overlap, *n* query size, *K* pathway size, and *N* background size.

Terms were processed in ascending FDR order, with ties resolved by decreasing enrichment factor. A term was assigned to an existing representative when shared candidate genes comprised at least 70% of the smaller matched set. FDR *<* 0.05 defined significance. For display, up to ten representative terms per candidate set were retained at ≤FDR 0.10, requiring at least three candidate genes. Their union formed the heatmap rows, grouped by the candidate set with the greatest enrichment factor and ordered by decreasing enrichment factor within each group. Entries failing display criteria were grey.

#### Day-10 reprogramming outcome and AUROC analysis

Reprogramming outcome was assessed in each donor’s D10 library using six endogenous pluripotency markers: *NANOG, LIN28A, DNMT3B, DPPA4, ZFP42*, and *L1TD1. POU5F1* and *SOX2* were excluded because their transcripts also derive from the delivery vector. A marker was considered detected when its expression was greater than zero. Cells detecting at least two markers were classified as marker-positive and cells detecting none as marker-negative; cells detecting one marker were excluded from ROC analyses.

Individual genes were scored by log-normalized expression, and the ISG score was the mean log-normalized expression of *ISG15, IFIT1, IFIT3*, and *IRF7*. AUROC treated marker-negative cells as the positive class. Confidence intervals for *DDX58* and the ISG score were obtained by class-stratified cell bootstrap within each donor (Table S6).

#### ISG expression and reprogramming outcome

This analysis included all D10 cells. Cells detecting at least two endogenous pluripotency markers were classified as marker-positive, and all other cells, including one-marker cells, as marker-negative. Within each donor, cells with an ISG score of zero formed one stratum, and cells with positive scores were divided into seven quantile strata. For each stratum, we calculated the mean ISG score and the marker-positive fraction with a Wilson 95% confidence interval. Because marker detection and ISG scores both depend on per-cell sequencing depth, donor-specific logistic regression modeled marker-positive status as a function of ISG score and log_10_ total UMI count. Fitted curves were evaluated at the donor’s median depth, and the exponentiated ISG coefficient gave the odds ratio per unit increase in ISG score. A sensitivity analysis replaced total UMI count with detected-gene count (Table S6).

#### Single-cell characterization of OSKM-induced intestinal partial reprogramming

GSE23717contains intestinal epithelial cells from doxycycline-treated (Dox) and untreated control (Cont) mice. Cell types were annotated with canonical markers from Kim et al.^54^, and YAP and fetal signature scores used gene sets from the same study. Partial-reprogramming and pluripotency signatures from Lu et al.^25^ were scored as described above, and Dox-minus-Cont differences in mean module score were calculated for each cell type with at least 50 cells in Dox. Slingshot trajectories toward revSC and aVEC were rooted at each candidate epithelial type. Developmental potential was scored with CytoTRACE2 v1.1.0, and for each candidate source, Dox and Cont scores were compared with a one-sided Wilcoxon rank-sum test (Dox *>* Cont).

#### Candidate-source prioritization in GSE237173

Control cells of each candidate source were paired with each target population (revSC or aVEC). Among Slingshot lineages rooted at that candidate, we selected the lineage containing the most target cells and excluded source cells outside it. Source populations were downsampled to 64 cells, the size of the smallest eligible population (TA). Each candidate–target pair was trained five times with independent subsamples and seeds, and rankings were merged by mean rank percentile.

Each candidate was evaluated by AUROC recovery of curated reprogramming regulators and chemical-cocktail targets within the 3,000-gene universe, excluding delivered factors. Candidates were compared by paired bootstrap resampling: reference and non-reference genes were resampled separately, and the same resample was applied to every candidate. We report the fraction of draws in which one candidate’s AUROC exceeded another’s. Recovery was also recomputed after removing annotated targets (TRRUST) (Table S7).

#### Expression-based assessment of the candidate sources

Expression-based assessments were performed for CBC, TA, EC, and PC, matching the FateDriver comparison, using control cells of the four candidates and target cells. For residual identity, candidate-specific markers were ranked against the other control types by a Wilcoxon rank-sum test, and the top 30 markers per candidate were scored in target cells with `scanpy.tl.score genes`. For label transfer, a *k* = 5 nearest-neighbor classifier trained on control cells in the Harmony-corrected embedding^68^ assigned candidate labels to target cells, and the assigned fraction was calculated for each candidate.

Both assessments used same number of control cells per candidate, redrawn 200 times, and intervals were the 2.5th and 97.5th percentiles across draws. The top candidate was tested against 200 candidate-label permutations, using the maximum across candidates as a family-wise null. As a positive control, each candidate was also compared with its nearest other control type. We also calculated candidate-wise differences in assigned fractions between revSC and aVEC (Table S7).

#### Lineage-specific differential expression and pathway enrichment

For each predicted source(TA or EC), three Slingshot lineages were compared using tradeSeq v1.20.0^70^: the lineage toward the partially reprogrammed target (L1; TA-to-revSC or EC-to-aVEC) and two alternative lineages ending in PC (L2) or CBC (L3). Lineages ending in EE or TC were excluded because these cell types comprised fewer than 50 cells. L1 was truncated at the maximum pseudo-time of the target cell type. A generalized additive model with three knots was fitted to highly variable genes (`fitGAM`), and endpoint differences were tested with `diffEndTest`. Genes with Benjamini–Hochberg FDR *<* 0.01 were retained, excluding those with |logFC|≥ 20. Genes with |logFC| *>* 1 in the same direction against both L2 and L3 were defined as lineage-specific upregulated or downregulated genes. KEGG enrichment was performed with `enrichKEGG` in clusterProfiler v4.14.6; pathways with FDR *<* 0.05 were considered significant.

### Quantification and statistical analysis

Unless otherwise specified, bootstrap analyses used 2,000 resamples with replacement and percentile 95% intervals from the 2.5th and 97.5th percentiles. Sampling units, stratification, and paired comparisons are defined for each analysis above. The expression-based origin assessment used repeated equal-size cell subsampling; ISG-group proportions used Wilson intervals. RNG seed 0 was used for delivered-factor rank bootstrap, source-state UMI downsampling, delivered-factor module scoring, and CellRank module scoring. Reference-recovery bootstrap initialized seed 0 for each dataset; chemical-target bootstrap initialized it for each method and target set. Day-10 AUROC bootstrap used seed 20260727. Values below the reporting cutoff were written as *p <* 1× 10^*−*300^. Analyses used Python and R versions listed in the key resources table.

### Additional resources

This study did not generate additional resources.

## References

1. Takahashi, K., and Yamanaka, S. (2016). A decade of transcription factor-mediated re-programming to pluripotency. Nature Reviews. Molecular Cell Biology 17, 183–193. doi: 10.1038/nrm.2016.8.

2. Cherry, A.B.C., and Daley, G.Q. (2013). Reprogrammed cells for disease modeling and regenerative medicine. Annual Review of Medicine 64, 277–290. doi: 10.1146/annurev-med-050311-163324.

3. Brumbaugh, J., Di Stefano, B., and Hochedlinger, K. (2019). Reprogramming: identifying the mechanisms that safeguard cell identity. Development (Cambridge, England) 146, dev182170. URL: https://pmc.ncbi.nlm.nih.gov/articles/PMC6918747/. doi: 10.1242/dev.182170.

4. Meir, R., Schwartz, G., Adam, M., Lapidot, A.A., Cain, A., Green, G.S., Menon, V., Bennett, D.A., De Jager, P.L., and Habib, N. (2025). Early neuronal reprogramming and cell cycle reentry shape Alzheimer’s disease progression. bioRxiv: The Preprint Server for Biology. doi: 10.1101/2025.06.04.653670 iSSN: 2692-8205.

5. Amo-Shiinoki, K., Tanabe, K., Hoshii, Y., Matsui, H., Harano, R., Fukuda, T., Takeuchi, T., Bouchi, R., Takagi, T., Hatanaka, M., Takeda, K., Okuya, S., Nishimura, W., Kudo, A., Tanaka, S., Tanabe, M., Akashi, T., Yamada, T., Ogawa, Y., Ikeda, E., Nagano, H., and Tanizawa, Y. (2021). Islet cell dedifferentiation is a pathologic mechanism of long-standing progression of type 2 diabetes. JCI insight 6, e143791, 143791. doi: 10.1172/jci.insight.143791.

6. Yuan, S., Norgard, R.J., and Stanger, B.Z. (2019). Cellular Plasticity in Cancer. Cancer Discovery 9, 837–851. doi: 10.1158/2159-8290.CD-19-0015.

7. de Lázaro, I., Cossu, G., and Kostarelos, K. (2017). Transient transcription factor (OSKM) expression is key towards clinical translation of in vivo cell reprogramming. EMBO molecular medicine 9, 733–736. doi: 10.15252/emmm.201707650.

8. Núñez-Quintela, V., Li, H., and Collado, M. (2026). Cellular reprogramming beyond pluripotency. Trends in Molecular Medicine pp. S1471–4914(26)00011–0. doi: 10.1016/j.molmed.2026.01.007.

9. Ieda, M., Fu, J.D., Delgado-Olguin, P., Vedantham, V., Hayashi, Y., Bruneau, B.G., and Srivastava, D. (2010). Direct reprogramming of fibroblasts into functional cardiomyocytes by defined factors. Cell 142, 375–386. doi: 10.1016/j.cell.2010.07.002. PMID: 20691899.

10. Huyghe, A., Furlan, G., Schroeder, J., Cascales, E., Trajkova, A., Ruel, M., Stüder, F., Larcombe, M., Bo Yang Sun, Y., Mugnier, F., De Matteo, L., Baygin, A., Wang, J., Yu, Y., Rama, N., Gibert, B., Kielbassa, J., Tonon, L., Wajda, P., Gadot, N., Brevet, M., Siouda, M., Mulligan, P., Dante, R., Liu, P., Gronemeyer, H., Mendoza-Parra, M., Polo, J.M., and Lavial, F. (2022). Comparative roadmaps of reprogramming and oncogenic transformation identify bcl11b and atoh8 as broad regulators of cellular plasticity. Nature Cell Biology 24, 1350–1363. doi: 10.1038/s41556-022-00986-w. PMID: 36075976; source dataset GSE181451.

11. Cheloufi, S., Elling, U., Hopfgartner, B., Jung, Y.L., Murn, J., Ninova, M., Hubmann, M., Badeaux, A.I., Euong Ang, C., Tenen, D., Wesche, D.J., Abazova, N., Hogue, M., Tasdemir, N., Brumbaugh, J., Rathert, P., Jude, J., Ferrari, F., Blanco, A., Fellner, M., Wenzel, D., Zinner, M., Vidal, S.E., Bell, O., Stadtfeld, M., Chang, H.Y., Almouzni, G., Lowe, S.W., Rinn, J., Wernig, M., Aravin, A., Shi, Y., Park, P.J., Penninger, J.M., Zuber, J., and Hochedlinger, K. (2015). The histone chaperone CAF-1 safeguards somatic cell identity. Nature 528, 218– URL: https://www.nature.com/articles/nature15749. doi: 10.1038/nature15749.

12. Huang, X., Balmer, S., Lyu, C., Xiang, Y., Malik, V., Wang, H., Zhang, Y., Cai, B., Xie, W., Hadjantonakis, A.K., Zhou, H., and Wang, J. (2024). ZFP281 controls transcriptional and epigenetic changes promoting mouse pluripotent state transitions via DNMT3 and TET1. Developmental Cell 59, 465–481.e6. URL:https://www.sciencedirect.com/science/article/pii/S1534580723006986. doi: 10.1016/j.devcel.2023.12.018.

13. Treutlein, B., Lee, Q.Y., Camp, J.G., Mall, M., Koh, W., Shariati, S.A.M., Sim, S., Neff, N.F., Skotheim, J.M., Wernig, M., and Quake, S.R. (2016). Dissecting direct reprogramming from fibroblast to neuron using single-cell RNA-seq. Nature 534, 391–395. URL: https://escholarship.org/uc/item/3wg3m7km. doi: 10.1038/nature18323.

14. Schiebinger, G., Shu, J., Tabaka, M., Cleary, B., Subramanian, V., Solomon, A., Gould, J., Liu, S., Lin, S., Berube, P., Lee, L., Chen, J., Brumbaugh, J., Rigollet, P., Hochedlinger, K., Jaenisch, R., Regev, A., and Lander, E.S. (2019). Optimal-Transport Analysis of Single-Cell Gene Expression Identifies Developmental Trajectories in Reprogramming. Cell 176, 928–943.e22. URL: https://www.sciencedirect.com/science/article/pii/S009286741930039X. doi: 10.1016/j.cell.2019.01.006.

15. Vainorius, G., Novatchkova, M., Michlits, G., Baar, J.C., Raupach, C., Lee, J., Yelagandula, R., Wernig, M., and Elling, U. (2023). Ascl1 and Ngn2 convert mouse embryonic stem cells to neurons via functionally distinct paths. Nature Communications 14, 5341. URL: https://www.nature.com/articles/s41467-023-40803-y. doi: 10.1038/s41467-023-40803-y.

16. Shu, J. (2025). When cellular reprogramming meets AI: towards de novo cell design. Nature Reviews. Genetics 26, 585. doi: 10.1038/s41576-025-00878-3.

17. Capponi, S., and Wang, S. (2024). AI in cellular engineering and reprogramming. Biophysical Journal 123, 2658–2670. URL: https://pmc.ncbi.nlm.nih.gov/articles/PMC11393708/. doi: 10.1016/j.bpj.2024.04.001.

18. Wang, P. et al. (2023). Deciphering driver regulators of cell fate decisions from single-cell transcriptomics data with cefcon. Nature Communications 14, 8459.

19. Martini, P., Hartebrodt, A., de Almeida, G.P., Hackstein, C.P., Zehn, D., and Blumenthal, D.B. (2025). Switchtfi: identifying transcription factors driving cell differentiation. Genome Biology 26, 410. doi: 10.1186/s13059-025-03876-0. PMID: 41331466.

20. Kim, J. et al. (2021). Tenet: gene network reconstruction using transfer entropy reveals key regulatory factors from single cell transcriptomic data. Nucleic Acids Research 49, e1.

21. Sadria, M., Layton, A., Goyal, S., and Bader, G.D. (2024). Fatecode enables cell fate regulator prediction using classification-supervised autoencoder perturbation. Cell Reports Methods 4, 100819. doi: 10.1016/j.crmeth.2024.100819.

22. Kiryo, R., Niu, G., du Plessis, M.C., and Sugiyama, M. (2017). Positive-unlabeled learning with non-negative risk estimator. In Advances in Neural Information Processing Systems vol. 30. pp. 1675–1685.

23. Bekker, J., and Davis, J. (2020). Learning from positive and unlabeled data: a survey. Machine Learning 109, 719–760. doi: 10.1007/s10994-020-05877-5.

24. Browaeys, R., Saelens, W., and Saeys, Y. (2020). Nichenet: modeling intercellular communication by linking ligands to target genes. Nature Methods 17, 159–162.

25. Lu, J.Y., Tu, W.B., Li, R., Weng, M., Sanketi, B.D., Yuan, B., Reddy, P., Rodriguez Esteban, C., and Izpisua Belmonte, J.C. (2025). Prevalent mesenchymal drift in aging and disease is reversed by partial reprogramming. Cell 188, 5895–5911.e17. doi: 10.1016/j.cell.2025.031. PMID: 40816266; source dataset GSE297234.

26. Liu, Z., Wang, L., Welch, J.D., Ma, H., Zhou, Y., Vaseghi, H.R., Yu, S., Wall, J.B., Alimohamadi, S., Zheng, M., Yin, C., Shen, W., Prins, J.F., Liu, J., and Qian, L. (2017). Single-cell transcriptomics reconstructs fate conversion from fibroblast to cardiomyocyte. Nature 551, 100–104. doi: 10.1038/nature24454. PMID: 29072293; source dataset GSE98567.

27. Ebrahimi, B. (2015). Reprogramming barriers and enhancers: strategies to enhance the efficiency and kinetics of induced pluripotency. Cell Regeneration 4, 10. doi: 10.1186/s13619-015-0024-9. PMID: 26566431; PMCID: PMC4642739; gene sets taken from Table 1.

28. Liu, M., Liu, J., Zhang, T., and Wang, L. (2022). Direct cardiac reprogramming: Toward the era of multi-omics analysis. Cell Insight 1, 100058. doi: 10.1016/j.cellin.2022.100058. PMID: 37193352.

29. Cui, Y., Zheng, Y., Liu, X., Yan, L., Fan, X., Yong, J., Hu, Y., Dong, J., Li, Q., Wu, X., Gao, S., Li, J., Ren, G., Wen, L., Tang, F., and Qiao, J. (2019). Single-cell transcriptome analysis maps the developmental track of the human heart. Cell Reports 26, 1934–1950.e5. doi: 10.1016/j.celrep.2019.01.079.

30. Cao, J., O’Day, D.R., Pliner, H.A., Kingsley, P.D., Deng, M., Daza, R.M., Zager, M.A., Aldinger, K.A., Blecher-Gonen, R., Zhang, F., Spielmann, M., Palis, J., Doherty, D., Steemers, F.J., Glass, I.A., Trapnell, C., and Shendure, J. (2020). A human cell atlas of fetal gene expression. Science 370, eaba7721. doi: 10.1126/science.aba7721.

31. Kanehisa, M., and Goto, S. (2000). Kegg: Kyoto encyclopedia of genes and genomes. Nucleic Acids Research 28, 27–30. doi: 10.1093/nar/28.1.27.

32. Ashburner, M., Ball, C.A., Blake, J.A., Botstein, D., Butler, H., Cherry, J.M., Davis, A.P., Dolinski, K., Dwight, S.S., Eppig, J.T., Harris, M.A., Hill, D.P., Issel-Tarver, L., Kasarskis, A., Lewis, S., Matese, J.C., Richardson, J.E., Ringwald, M., Rubin, G.M., and Sherlock, G. (2000). Gene ontology: tool for the unification of biology. Nature Genetics 25, 25–29. doi: 10.1038/75556.

33. Subramanian, A., Tamayo, P., Mootha, V.K., Mukherjee, S., Ebert, B.L., Gillette, M.A., Paulovich, A., Pomeroy, S.L., Golub, T.R., Lander, E.S., and Mesirov, J.P. (2005). Gene set enrichment analysis: a knowledge-based approach for interpreting genome-wide expression profiles. Proceedings of the National Academy of Sciences 102, 15545–15550. doi: 10.1073/pnas.0506580102.

34. Liberzon, A., Subramanian, A., Pinchback, R., Thorvaldsdóttir, H., Tamayo, P., and Mesirov, J.P. (2011). Molecular signatures database (msigdb) 3.0. Bioinformatics 27, 1739–1740. doi: 10.1093/bioinformatics/btr260.

35. Castanza, A.S., Recla, J.M., Eby, D., Thorvaldsdóttir, H., Bult, C.J., and Mesirov, J.P. (2023). Extending support for mouse data in the molecular signatures database (msigdb). Nature Methods 20, 1619–1620. doi: 10.1038/s41592-023-02014-7.

36. Hou, P., Li, Y., Zhang, X., Liu, C., Guan, J., Li, H., Zhao, T., Ye, J., Yang, W., Liu, K., Ge, J., Xu, J., Zhang, Q., Zhao, Y., and Deng, H. (2013). Pluripotent stem cells induced from mouse somatic cells by small-molecule compounds. Science 341, 651–654. doi: 10.1126/science.1239278. PMID: 23868920.

37. Zhao, Y., Zhao, T., Guan, J., Zhang, X., Fu, Y., Ye, J., Zhu, J., Meng, G., Ge, J., Yang, S., Cheng, L., Du, Y., Zhao, C., Wang, T., and Deng, H. (2015). A XEN-like state bridges somatic cells to pluripotency during chemical reprogramming. Cell 163, 1678–1691. doi: 10.1016/j.cell.2015.11.017. PMID: 26686652.

38. Guan, J., Wang, G., Wang, J., Zhang, Z., Fu, Y., Cheng, L., Meng, G., Lyu, Y., Zhu, J., Li, Y., Wang, Y., Liuyang, S., Liu, B., Yang, Z., and Deng, H. (2022). Chemical reprogramming of human somatic cells to pluripotent stem cells. Nature 605, 325–331. doi: 10.1038/s41586-022-04593-5. PMID: 35418683.

39. Liuyang, S., Wang, G., Wang, Y., He, H., Lyu, Y., Cheng, L., Yang, Z., Guan, J., Fu, Y., Zhu, J., Zhong, X., Sun, S., Li, C., Wang, J., and Deng, H. (2023). Highly efficient and rapid generation of human pluripotent stem cells by chemical reprogramming. Cell Stem Cell 30, 450–459.e9. doi: 10.1016/j.stem.2023.02.008. PMID: 36944335.

40. Wang, Y., Peng, F., Yang, Z., Cheng, L., Cao, J., Fu, X., He, H., Cai, R., Zeng, W., Dong, Y., Chen, G., Peng, G., Liuyang, S., Wang, G., and Deng, H. (2025). A rapid chemical reprogramming system to generate human pluripotent stem cells. Nature Chemical Biology 21, 1030–1038. doi: 10.1038/s41589-024-01799-8. PMID: 39753706.

41. Fu, Y., Huang, C., Xu, X., Gu, H., Ye, Y., Jiang, C., Qiu, Z., and Xie, X. (2015). Direct reprogramming of mouse fibroblasts into cardiomyocytes with chemical cocktails. Cell Research 25, 1013–1024. doi: 10.1038/cr.2015.99. PMID: 26292833.

42. Cao, N., Huang, Y., Zheng, J., Spencer, C.I., Zhang, Y., Fu, J.D., Nie, B., Xie, M., Zhang, M., Wang, H., Ma, T., Xu, T., Shi, G., Srivastava, D., and Ding, S. (2016). Conversion of human fibroblasts into functional cardiomyocytes by small molecules. Science 352, 1216–1220. doi: 10.1126/science.aaf1502. PMID: 27127239.

43. Park, G., Yoon, B.S., Kim, Y.S., Choi, S.C., Moon, J.H., Kwon, S., Hwang, J., Yun, W., Kim, J.H., Park, C.Y., Lim, D.S., Kim, Y.I., Oh, C.H., and You, S. (2015). Conversion of mouse fibroblasts into cardiomyocyte-like cells using small molecule treatments. Biomaterials 54, 201–212. doi: 10.1016/j.biomaterials.2015.02.029. PMID: 25907053.

44. Huang, C., Tu, W., Fu, Y., Wang, J., and Xie, X. (2018). Chemical-induced cardiac reprogramming in vivo. Cell Research 28, 686–689. doi: 10.1038/s41422-018-0036-4. PMID: 29670223.

45. Chang, D., Sun, C., Tian, X., Liu, H., Jia, Y., and Guo, Z. (2024). Regulation of cardiac fibroblasts reprogramming into cardiomyocyte-like cells with a cocktail of small molecule compounds. FEBS Open Bio 14, 983–1000. doi: 10.1002/2211-5463.13811. PMID: 38693086.

46. Turchaninova, E., Robustova, S., Kovalenko, S., Dzhabrailov, V., Dolgodvorova, A., Romanova, S., Zybin, D., Popov, M., Miftakhova, A., Frolova, S., Slotvitsky, M., Romanov, A., Agladze, K., and Tsvelaya, V. (2026). Refining fibroblast-to-cardiomyocyte transdifferentiation protocols to explore emergent self-organization in cardiac cultures. PLoS One 21, e0343415. doi: 10.1371/journal.pone.0343415.

47. Banito, A., Rashid, S.T., Acosta, J.C., Li, S., Pereira, C.F., Geti, I., Pinho, S., Silva, J.C., Azuara, V., Walsh, M., Vallier, L., and Gil, J. (2009). Senescence impairs successful reprogramming to pluripotent stem cells. Genes & Development 23, 2134–2139. doi: 10.1101/gad.1811609.

48. Li, H., Collado, M., Villasante, A., Strati, K., Ortega, S., Cañamero, M., Blasco, M.A., and Serrano, M. (2009). The ink4/arf locus is a barrier for ips cell reprogramming. Nature 460, 1136–1139. doi: 10.1038/nature08290.

49. Li, R., Liang, J., Ni, S., Zhou, T., Qing, X., Li, H., He, W., Chen, J., Li, F., Zhuang, Q., Qin, B., Xu, J., Li, W., Yang, J., Gan, Y., Qin, D., Feng, S., Song, H., Yang, D., Zhang, B., Zeng, L., Lai, L., Esteban, M.A., and Pei, D. (2010). A mesenchymal-to-epithelial transition initiates and is required for the nuclear reprogramming of mouse fibroblasts. Cell Stem Cell 7, 51–63. doi: 10.1016/j.stem.2010.04.014.

50. Lange, M., Bergen, V., Klein, M., Setty, M., Reuter, B., Bakhti, M., Lickert, H., Ansari, M., Schniering, J., Schiller, H.B., Pe’er, D., and Theis, F.J. (2022). Cellrank for directed single-cell fate mapping. Nature Methods 19, 159–170. doi: 10.1038/s41592-021-01346-6.

51. Andrews, P.W., and Gokhale, P.J. (2024). A short history of pluripotent stem cells markers. Stem Cell Reports 19, 1–10. doi: 10.1016/j.stemcr.2023.11.012. PMID: 38157849.

52. O’Malley, J., Skylaki, S., Iwabuchi, K.A., Chantzoura, E., Ruetz, T., Johnsson, A., Tomlinson, S.R., Linnarsson, S., and Kaji, K. (2013). High-resolution analysis with novel cell-surface markers identifies routes to ips cells. Nature 499, 88–91. doi: 10.1038/nature12243. PMID: 23728301.

53. Yoneyama, M., Kikuchi, M., Natsukawa, T., Shinobu, N., Imaizumi, T., Miyagishi, M., Taira, K., Akira, S., and Fujita, T. (2004). The rna helicase rig-i has an essential function in double-stranded rna-induced innate antiviral responses. Nature Immunology 5, 730–737. doi: 10.1038/ni1087.

54. Kim, J., Kim, S., Lee, S.Y., Jo, B.K., Oh, J.Y., Kwon, E.J., Kim, K.T., Adpaikar, A.A., Kim, E.J., Jung, H.S., Kim, H.R., Roe, J.S., Hong, C.P., Kim, J.K., Koo, B.K., and Cha, H.J. (2023). Partial in vivo reprogramming enables injury-free intestinal regeneration via autonomous ptgs1 induction. Science Advances 9, eadi8454. doi: 10.1126/sciadv.adi8454. PMID: 38000027; source dataset GSE237173.

55. Kang, M., Gulati, G.S., Brown, E.L., Qi, Z., Avagyan, S., Armenteros, J.J.A., Gleyzer, R., Zhang, W., Steen, C.B., D’Silva, J.P., Schwab, J., Clarke, M.F., Chaudhuri, A.A., and New-man, A.M. (2025). Improved reconstruction of single-cell developmental potential with CytoTRACE 2. Nature Methods 22, 2258–2263. doi: 10.1038/s41592-025-02857-2.

56. Samavarchi-Tehrani, P., Golipour, A., David, L., Sung, H.K., Beyer, T.A., Datti, A., Woltjen, K., Nagy, A., and Wrana, J.L. (2010). Functional genomics reveals a BMP-driven mesenchymal-to-epithelial transition in the initiation of somatic cell reprogramming. Cell Stem Cell 7, 64–77. doi: 10.1016/j.stem.2010.04.015. PMID: 20621051.

57. Ocampo, A., Reddy, P., Martinez-Redondo, P., Platero-Luengo, A., Hatanaka, F., Hishida, T., Li, M., Lam, D., Kurita, M., Beyret, E., Araoka, T., Vazquez-Ferrer, E., Donoso, D., Roman, J.L., Xu, J., Rodriguez Esteban, C., Nuñez, G., Nuñez Delicado, E., Campistol, J.M., Guillen, I., Guillen, P., and Izpisua Belmonte, J.C. (2016). In Vivo Amelioration of Age-Associated Hallmarks by Partial Reprogramming. Cell 167, 1719–1733.e12. doi: 10.1016/j.cell.2016.11.052.

58. Nishimura, K., Sano, M., Ohtaka, M., Furuta, B., Umemura, Y., Nakajima, Y., Ikehara, Y., Kobayashi, T., Segawa, H., Takayasu, S., Sato, H., Motomura, K., Uchida, E., Kanayasu-Toyoda, T., Asashima, M., Nakauchi, H., Yamaguchi, T., and Nakanishi, M. (2011). Development of defective and persistent Sendai virus vector: A unique gene delivery/expression system ideal for cell reprogramming. Journal of Biological Chemistry 286, 4760–4771. doi: 10.1074/jbc.M110.183780. PMID: 21138846.

59. Han, H., Cho, J.W., Lee, S., Yun, A., Kim, H., Bae, D., Yang, S., Kim, C.Y., Lee, M., Kim, E., Lee, S., Kang, B., Jeong, D., Kim, Y., Jeon, H.N., Jung, H., Nam, S., Chung, M., Kim, J.H., and Lee, I. (2018). TRRUST v2: an expanded reference database of human and mouse transcriptional regulatory interactions. Nucleic Acids Research 46, D380–D386. doi: 10.1093/nar/gkx1013.

60. Wolf, F.A., Angerer, P., and Theis, F.J. (2018). Scanpy: large-scale single-cell gene expression data analysis. Genome Biology 19, 15. doi: 10.1186/s13059-017-1382-0.

61. Van de Sande, B., Flerin, C., Davie, K. et al. (2020). A scalable scenic workflow for single-cell gene regulatory network analysis. Nature Protocols 15, 2247–2276.

62. Wu, T., Hu, E., Xu, S., Chen, M., Guo, P., Dai, Z., Feng, T., Zhou, L., Tang, W., Zhan, L., Fu, X., Liu, S., Bo, X., and Yu, G. (2021). clusterprofiler 4.0: A universal enrichment tool for interpreting omics data. The Innovation 2, 100141. doi: 10.1016/j.xinn.2021.100141.

63. Liberzon, A., Birger, C., Thorvaldsdóttir, H., Ghandi, M., Mesirov, J.P., and Tamayo, P. (2015). The molecular signatures database hallmark gene set collection. Cell Systems 1, 417–425. doi: 10.1016/j.cels.2015.12.004.

64. Street, K., Risso, D., Fletcher, R.B., Das, D., Ngai, J., Yosef, N., Purdom, E., and Dudoit, S. (2018). Slingshot: cell lineage and pseudotime inference for single-cell transcriptomics. BMC Genomics 19, 477. doi: 10.1186/s12864-018-4772-0.

65. Hao, Y., Stuart, T., Kowalski, M.H., Choudhary, S., Hoffman, P., Hartman, A., Srivastava, A., Molla, G., Madad, S., Fernandez-Granda, C., and Satija, R. (2024). Dictionary learning for integrative, multimodal and scalable single-cell analysis. Nature Biotechnology 42, 293–304. doi: 10.1038/s41587-023-01767-y.

66. Wolock, S.L., Lopez, R., and Klein, A.M. (2019). Scrublet: Computational identification of cell doublets in single-cell transcriptomic data. Cell Systems 8, 281–291. doi: 10.1016/j.cels.2018.11.005.

67. Germain, P.L., Lun, A., Garcia Meixide, C., Macnair, W., and Robinson, M.D. (2021). Doublet identification in single-cell sequencing data using scdblfinder. F1000Research 10, 979. doi: 10.12688/f1000research.73600.2.

68. Korsunsky, I., Millard, N., Fan, J., Slowikowski, K., Zhang, F., Wei, K., Baglaenko, Y., Brenner, M., Loh, P.r., and Raychaudhuri, S. (2019). Fast, sensitive and accurate integration of single-cell data with harmony. Nature Methods 16, 1289–1296. doi: 10.1038/s41592-019-0619-0.

69. Setty, M., Kiseliovas, V., Levine, J., Gayoso, A., Mazutis, L., and Pe’er, D. (2019). Characterization of cell fate probabilities in single-cell data with palantir. Nature Biotechnology 37, 451–460. doi: 10.1038/s41587-019-0068-4.

70. Van den Berge, K., Roux de Bézieux, H., Street, K., Saelens, W., Cannoodt, R., Saeys, Y., Dudoit, S., and Clement, L. (2020). Trajectory-based differential expression analysis for single-cell sequencing data. Nature Communications 11, 1201. doi: 10.1038/s41467-020-14766-3.

71. Halko, N., Martinsson, P.G., and Tropp, J.A. (2011). Finding structure with randomness: probabilistic algorithms for constructing approximate matrix decompositions. SIAM Review 53, 217–288.

72. Kipf, T.N., and Welling, M. (2017). Semi-supervised classification with graph convolutional networks. In International Conference on Learning Representations (ICLR).

73. Lopez, R., Regier, J., Cole, M.B., Jordan, M.I., and Yosef, N. (2018). Deep generative modeling for single-cell transcriptomics. Nature Methods 15, 1053–1058.

74. Lee, J., Lee, Y., Kim, J., Kosiorek, A.R., Choi, S., and Teh, Y.W. (2019). Set transformer: a framework for attention-based permutation-invariant neural networks. In International Conference on Machine Learning (ICML). pp. 3744–3753.

75. Vaswani, A., Shazeer, N., Parmar, N., Uszkoreit, J., Jones, L., Gomez, A.N., Kaiser, Ł., and Polosukhin, I. (2017). Attention is all you need. In Advances in Neural Information Processing Systems (NeurIPS) vol. 30. pp. 5998–6008.

76. Kim, S., Chen, J., Cheng, T., Gindulyte, A., He, J., He, S., Li, Q., Shoemaker, B.A., Thiessen, P.A., Yu, B., Zaslavsky, L., Zhang, J., and Bolton, E.E. (2025). PubChem 2025 update. Nucleic Acids Research 53, D1516–D1525. doi: 10.1093/nar/gkae1059. PMID: 39558165.

77. Knox, C., Wilson, M., Klinger, C.M., Franklin, M., Oler, E., Wilson, A., Pon, A., Cox, J., Chin, N.E.L., Strawbridge, S.A., Garcia-Patino, M., Kruger, R., Sivakumaran, A., Sanford, S., and Wishart, D.S. (2024). DrugBank 6.0: the DrugBank knowledgebase for 2024. Nucleic Acids Research 52, D1265–D1275. doi: 10.1093/nar/gkad976. PMID: 37953279.

78. Gilson, M.K., Liu, T., Baitaluk, M., Nicola, G., Hwang, L., and Chong, J. (2016). BindingDB in 2015: a public database for medicinal chemistry, computational chemistry and systems pharmacology. Nucleic Acids Research 44, D1045–D1053. doi: 10.1093/nar/gkv1072. PMID: 26481362.

79. The UniProt Consortium (2025). UniProt: the Universal Protein Knowledgebase in 2025. Nucleic Acids Research 53, D609–D617. doi: 10.1093/nar/gkae1010. PMID: 39552041.

